# A high-throughput CRISPR interference screen for dissecting functional regulators of GPCR/cAMP signaling

**DOI:** 10.1101/2020.09.02.279679

**Authors:** Khairunnisa Mentari Semesta, Ruilin Tian, Martin Kampmann, Mark von Zastrow, Nikoleta G. Tsvetanova

## Abstract

G protein-coupled receptors (GPCRs) allow cells to respond to chemical and sensory stimuli through generation of second messengers, such as cyclic AMP (cAMP), which in turn mediate a myriad of processes, including cell survival, proliferation, and differentiation. In order to gain deeper insights into the complex biology and physiology of these key cellular pathways, it is critical to be able to globally map the molecular factors that shape cascade function. Yet, to this date, efforts to systematically identify regulators of GPCR/cAMP signaling have been lacking. Here, we combined genome-wide screening based on CRISPR interference with a novel sortable transcriptional reporter that provides robust readout for cAMP signaling, and carried out a functional screen for regulators of the pathway. Due to the sortable nature of the platform, we were able to assay regulators with strong and weaker phenotypes by analyzing sgRNA distribution among three fractions with distinct reporter expression. We identified 45 regulators with strong and 50 regulators with weaker phenotypes not previously known to be involved in cAMP signaling. In follow-up experiments, we validated the functional effects of seven newly discovered mediators (*NUP93, PRIM1, RUVBL1, PKMYT1, TP53, SF3A2*, and *HRAS*), and showed that they control distinct steps of the pathway. Thus, our study provides proof of principle that the screening platform can be applied successfully to identify bona fide regulators of GPCR/second messenger cascades in an unbiased and high-throughput manner, and illuminates the remarkable functional diversity among GPCR regulators.

**Author summary:** Cells sense and respond to changes in their surrounding environment through G protein-coupled receptors (GPCRs) and their associated cascades. The proper function of these pathways is essential to human physiology, and GPCRs have become a prime target for drug development for a range of human diseases. Therefore, it is of utmost importance to be able to map how these pathways operate to enable cells to fine-tune their responsiveness. Here, we describe a screening approach that we have devised to systematically identify regulators of GPCR function. We have developed a sortable reporter system and coupled that with silencing of genes across the entire human genome in order to uncover a range of novel mediators of GPCR activity. We characterize a few of these new regulators and show that they function at different steps of the cascade. Therefore, this study serves as proof of principle for the new screening platform. We envision that the approach can be used to dissect additional dimensions of GPCR function, including regulators of drug-specific responses, functional characterization of receptor features, and identification of novel drugs, and thus advance a genome-scale understanding of these critical pathways.

## Introduction

The GPCR/cAMP cascade is a key signaling axis coordinating the ability of cells to respond to fluctuations in their environment. The pathway is initiated by ligand binding to the transmembrane receptor, which leads to the stimulation of adenylyl cyclase enzymes (ACs) by the dissociated G protein subunit, Gαs, and to subsequent cAMP production. cAMP binds to intracellular effectors, including its main effector, protein kinase A (PKA), and triggers substrate phosphorylation, and ultimately gene expression reprogramming through phosphorylation and activation of the nuclear cAMP-response binding protein (CREB). Because GPCRs are the largest and most versatile family of mammalian membrane-bound receptors and cAMP is a ubiquitous diffusible second messenger, this cascade mediates most of the essential aspects of human physiology and is a prime target for drug development for a range of human diseases.^1^ Therefore, the ability to systematically dissect the factors that regulate the function and outcomes of these pathways can illuminate the molecular underpinnings of complex physiology and pathophysiology, and identify novel candidates for selective and efficient therapeutic intervention.

Recent technological advances using CRISPR/Cas9-based methods have revolutionized the field of functional genomics and enabled large-scale unbiased screens to probe the regulation of signaling cascades. While these approaches have been successfully implemented to characterize a number of important mammalian pathways^2^, to our best knowledge there have been no efforts to date to systematically identify genes mediating GPCR/cAMP signaling. In large part, this can be attributed to the lack of robust selectable phenotypic readouts that such pooled screening methods necessitate. To bridge this gap, we optimized a fluorescent transcriptional reporter for cAMP signaling with low background and high dynamic range, and expressed it in a CRISPR-compatible cell line. Next, we combined the reporter with small guide RNA (sgRNA) library transduction and fluorescence-activated cell sorting to enable a high-throughput genome-scale CRISPR interference (CRISPRi)-based screen for modulators of GPCR signaling. We provide the proof of principle that this unbiased platform can identify bona fide novel regulators by applying it to dissect a prototypical GPCR/cAMP cascade, the beta2-adrenergic receptor (β2-AR)/cAMP pathway, a central mediator of cardiovascular and pulmonary physiology.^3,4^ Through pairwise comparisons between sorted fractions with distinct reporter expression, we identified 46 hits with strong and 50 hits with weaker phenotypes. Surprisingly, with the exception of the *ADRB2* gene encoding the β2-AR, the remaining 95 factors have not been previously linked to these cascades. We functionally validated seven of the newly discovered regulators (*PKMYT1, TP53, NUP93, PRIM1, RUVBL1, HRAS*, and *SF3A2*) by cloning individual sgRNAs and re-testing their effects on reporter expression. Finally, we selected two of the novel hits, the nucleoporin-encoding gene *NUP93* and the ER/Golgi kinase encoding *PKMYT1*, for in-depth characterization and showed that these factors impacts cAMP signaling and transcriptional responses by regulating distinct steps of the GPCR cascade. In conclusion, the high-throughput screening paradigm described here provides a novel platform for unbiased dissection of regulators of receptor function, and pinpoints target genes for future manipulation of the pathway and its signaling consequences.

## Results

### Generation of a fluorescent CREB reporter cell line for analysis of GPCR/cAMP signaling

The CRISPR interference (CRISPRi) technology relies on transcriptional repression and has been applied successfully in pooled screens to assess the functional impacts of silencing specific genes.^5^ We reasoned that optimizing a sortable readout for cAMP that has low background and high dynamic range would enable us to reliably capture a range of phenotypes in a pooled CRISPRi-based screen. Therefore, we took advantage of a CREB reporter, in which cAMP response elements (CREs) drive the expression of the fluorescent protein GFP downstream of a destabilizing domain (DD) (Fig. 1a). This degron element targets the GFP for proteasomal degradation under normal conditions but can be selectively stabilized with a cell permeable compound, Shield-1^6^, and thus can boost the dynamic range of the system. To test the performance of the sensor, we began by transiently transfecting HEK293 cells with the construct. HEK293 cells express functional β2-AR^7^ and are a useful system to study signaling effects at native receptor levels. We stimulated cAMP production in each of two ways: 1) through activation of the β2-AR/Gαs pathway with saturating doses of the full agonist isoproterenol, and 2) by direct AC activation with the drug forskolin (Fig. S1a). Using flow cytometry as readout, we saw an increase in GFP fluorescence after drug treatment only when the degron was stabilized by the simultaneous addition of Shield-1 (Fig. S1b), which confirmed that the reporter undergoes efficient proteasomal turnover in the absence of the chemical stabilizer. However, the induction in reporter expression in transiently transfected cells was modest-only ~2-fold with both isoproterenol and forskolin. In order to improve the dynamic range of the system, we next constructed a clonal reporter cell line hereafter called CRE-GFP/CRISPR. For that purpose, we cloned the sensor into a lentiviral vector, with which we transduced HEK293 cells, in combination with a catalytically dead Cas9-BFP enzyme fused to KRAB (a transcriptional repressor domain), and selected high reporter-expressing CRE-GFP/CRISPR cells (Fig. 1a).

**Figure 1.**
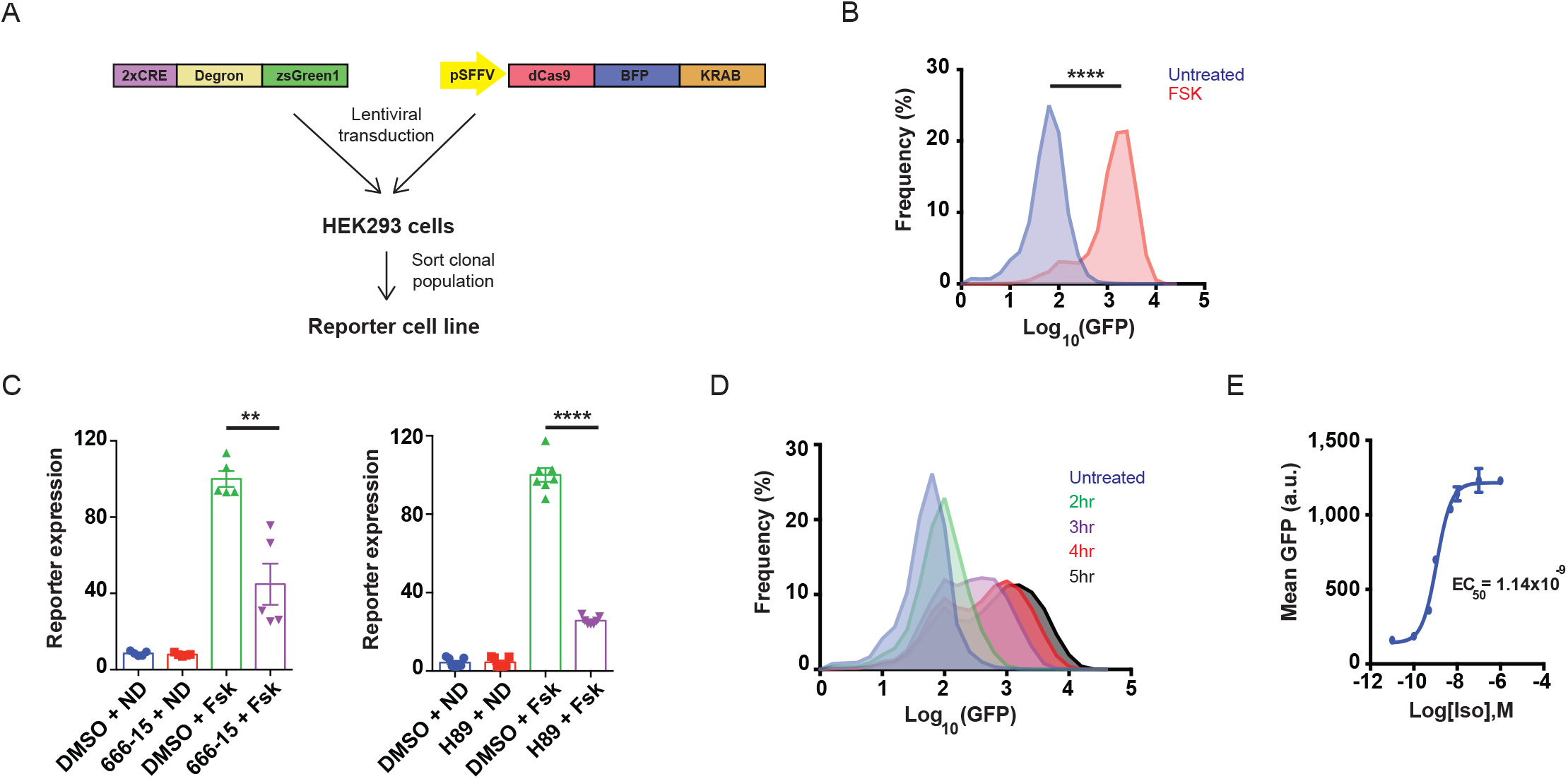
A robust CREB transcriptional reporter for GPCR/cAMP activity. **(A)** Schematic representation of the CREB reporter used in the CRISPR-based screen. Two cAMP response elements were fused to a ProteoTuner destabilizing domain (degron) followed by a green fluorescent protein, and inserted into a lentiviral vector. A clonal cell line is generated by transducing HEK293 cells with the reporter and dCas9-BFP-KRAB, sorting individual cells, and growing and verifying clonal lines for high reporter expression and efficient dCas9-dependent gene silencing. **(B)** The CREB reporter responds robustly to direct stimulation of adenylyl cyclase/cAMP signaling with forskolin. Reporter cells were treated with 10 μM forskolin (FSK) or DMSO (vehicle), and 1 μM Shield-1 was added simultaneously to stabilize the degron domain. After 4 h, reporter expression was analyzed by flow cytometry. Data from n = 4 per condition. **(C)** The CREB and PKA inhibitors, 666-15 and H89, respectively, significantly diminish FSK-induced accumulation of the reporter. Cells were pre-incubated with 100 nM 666-15 or 10 μM H89 versus DMSO (vehicle) for 30 min, then treated with 10 μM FSK in the presence of 1 μM Shield-1. After 4 h, reporter accumulation was analyzed by flow cytometry. Data plotted are from n = 5 for 666-15 and n = 7 for H89. **(D)** Timecourse for β2-AR-dependent reporter induction. Reporter cells were treated with 1 μM isoproterenol / 1 μM Shield-1 for indicated times or treated with 1 μM Shield-1 alone (no isoproterenol) for 5 h (“untreated”), and reporter expression was analyzed by flow cytometry. Data from n = 5. **(E)** Dose-response curve for isoproterenol-dependent reporter expression. Reporter cells were treated with indicated doses of isoproterenol and 1 μM Shield-1 for 4 h, and reporter expression was analyzed by flow cytometry. Data plotted are means of GFP expression, n = 3 per condition. EC_50_ curve-fitting was performed using Prism6 GraphPad software. In each flow cytometry experiment, 10,000 cells total were analyzed and gated for singlets. Error bars = ± s.e.m. **** = *p* ≤ 0.0001; ** = *p* ≤ 0.01 by unpaired two-tailed Student’s t-test.

To validate the suitability of the cell line for identification of functional regulators of GPCR signaling, we first measured reporter activity in response to cAMP production. Direct AC stimulation with forskolin led to robust accumulation of GFP compared to vehicle-treated control (> 20-fold increase, *p* ≤ 1.0×10^-4^ by unpaired Student’s t-test) (Fig. 1b). Notably, reporter activity was dependent on cAMP and CREB, because the magnitude of the response was markedly reduced upon acute blockade of either PKA, which regulates CREB activation, or CREB itself with cell-permeable small-molecule inhibitors (Fig. 1c) in agreement with previous reports.^8–10^ Next, we tested reporter activity upon β2-AR-dependent stimulation of cAMP signaling. We treated CRE-GFP/CRISPR cells with saturating doses of isoproterenol for varying intervals (0.5 - 5 h), and assessed GFP expression levels by flow cytometry. We observed accumulation of the reporter as early as 2 h after adrenoceptor activation that became more pronounced with longer treatments (Fig. 1d). At 4 h after isoproterenol addition, reporter levels were elevated > 8-fold relative to untreated cells, and this increase in GFP fluorescence was dose-dependent and saturable (EC_50_ ~ 1 nM isoproterenol, max ≥ 10 nM isoproterenol) (Fig. 1d-e, S1c). Taken together, these results demonstrate that CRE-GFP/CRISPR cells generate a robust and dynamic fluorescent readout in response to cAMP signaling from direct and GPCR-dependent stimulation of ACs. We chose a 4 h stimulation window for our future experiments, since this duration of treatment would meet practical considerations of a screen associated with time required for harvesting and sorting a large number of cells for analysis.

We next tested the efficiency of dCas9/CRISPR gene silencing in our cells. We began by targeting the *TFRC* gene, which encodes the ubiquitously expressed receptor for iron uptake, transferrin, as cell surface levels of transferrin can be easily and accurately quantified by antibody labeling and flow cytometry.^11^ We transduced CRE-GFP/CRISPR cells with non-targeting control (NTC) or *TFRC* sgRNAs and saw severe depletion of transferrin protein in *TFRC* sgRNA-expressing cells (Fig. S2a). Since our ultimate goal was to identify regulators of GPCR signaling using the β2-AR as a model, we next targeted the adrenoceptor-coding gene, *ADRB2*, and measured isoproterenol-dependent CRE-GFP accumulation by flow cytometry. Independent transduction with two different sgRNAs against *ADRB2* significantly blunted the transcriptional response to isoproterenol compared to NTCs (4-6-fold, *p* ≤ 1.0×10^-4^ by one-way ANOVA test) (Fig. S2b). In contrast, direct stimulation of CREB by addition of the cell-permeable cAMP analog, 8-bromo-cAMP, which bypasses the requirement for β2-AR activity, resulted in normal reporter induction in these cells (Fig. S2b). These results show that CRE-GFP/CRISPR cells are sensitive to CRISPRi-based perturbations of individual genes.

### The reporter cell line is compatible with CRISPR-based pooled screening

To determine the suitability of CRE-GFP/CRISPR for functional genomic screening, we next conducted a small-scale CRISPRi-based screen. For these experiments, we selected the kinase/phosphatase “H1” sub-genome library, which is one of the 7 CRISPRi libraries collectively targeting the vast majority of the annotated human genome (Table S1). H1 contains ~13,000 sgRNAs targeting 2,325 genes (5 sgRNAs per gene and 250 NTCs), among which-the *ADRB2*, making it ideal for testing the performance of our cell line in pooled screening. In addition, this sub-genome library contains sgRNAs targeting genes with redundant regulatory roles in the pathway, including the three GPCR kinases expressed in HEK293 cells, *GRK2, 5* and *6*, which act in concert to phosphorylate the β2-AR and promote its desensitization^12–14^, and the catalytic and regulatory subunits of PKA, *PRKAC* and *PRKAR*.

We packaged the H1 sgRNAs into lentiviral particles and transduced ~20 million CRE-GFP/CRISPR cells at low multiplicity of infection to ensure < 1 sgRNA/cell and sufficient coverage of the library in order to avoid depletion of sgRNAs targeting essential genes during the selection period. Clones with successful sgRNA integration were selected with puromycin and expanded for 1 week total following established protocols^15,16^, then treated with saturating doses of isoproterenol (1 μM), and sorted into three fractions (“low”, “medium” and “high”) based on GFP expression (Fig. 2a). Each sorted fraction contained ≥ 100-fold cell coverage of library elements. Gene enrichment scores (epsilon) and *p*-values were calculated as previously described.^17^ Of all ~2,300 targeted genes, *ADRB2* depletion gave rise to the most extreme downregulation of GFP reporter expression (> 14-fold sgRNA enrichment in “low” vs “high”, >5-fold in “medium” vs “high” and >2.5-fold in “medium” vs “low”) (Fig. 2b). Even though only the 3 sgRNAs with the most extreme phenotype were considered when calculating an epsilon value (see “Materials and Methods”), 4/5 sgRNAs against *ADRB2* exhibited > 4-fold enrichment in the “low” vs “high” fraction (Fig. 2c). These results are consistent with the severe blockade in transcriptional reporter activity in CRE-GFP/CRISPR cells that we observed upon *ADRB2* knockdown with individual sgRNAs (Fig. S2b). At the same time, the phenotype from CRISPRi-based depletion of different PKA subunits was much more subtle-only sgRNAs against the regulatory subunit genes RIb and RIIa showed enrichment in “low” versus “high” fraction (~1.5-fold, *p* = 1.2×10^-2^ for RIb and 8.2×10^-2^ for RIIa), while depletion of neither of the two catalytic subunit-encoding genes, Ca and Cb, had a significant effect (Fig. 2b). Similarly, we observed a modest enrichment (~1.3-fold) of sgRNAs in the “high” versus “low” fraction only for the negative regulator GRK2, but not GRKs 5 and 6 (Fig. 2b), a result that is consistent with previous studies showing functional redundancy of GRK5/6 in GPCR inactivation.^18,19^ These data confirm that CRE-GFP/CRISPR is suitable for detecting known regulators of the cAMP/CREB pathway in a pooled screen. However, they also highlight the likelihood for genes with redundant roles in the pathway to yield modest phenotypes.

**Figure 2.**
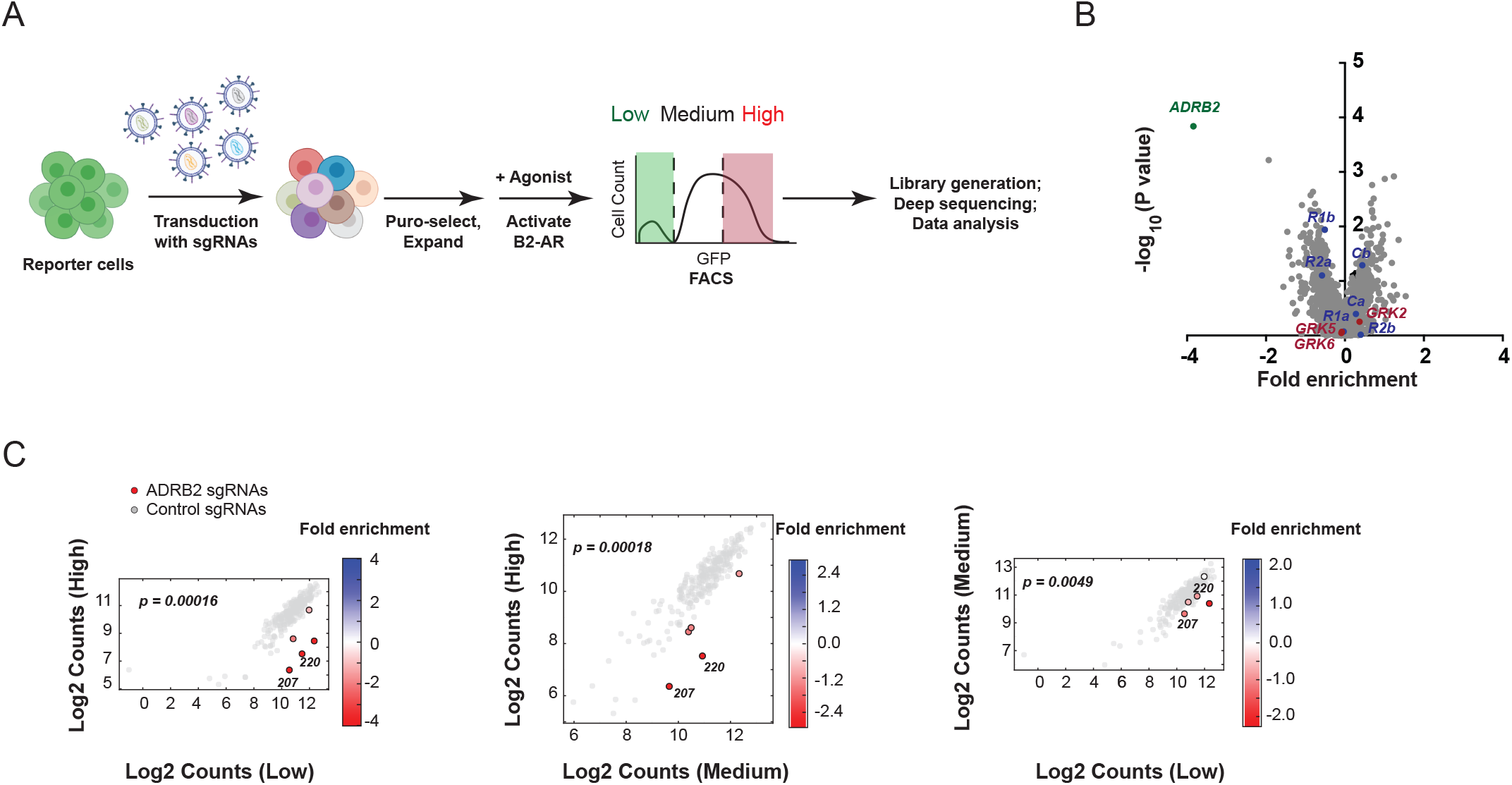
The reporter cell line is compatible with pooled functional genomic screening. **(A)** Schematic representation of the CRISPR interference-based genomic screen. Reporter cells were transduced with sgRNA libraries for 1 week, β2-AR signaling was activated with 1 μM isoproterenol/1 μM Shield-1 for 4 h, and edited cells were sorted based on differential expression of the CREB reporter into “low”, “medium” or “high” bins. Genomic DNA was extracted from each fraction, amplified, barcoded and subjected to deep sequencing and statistical analysis of sgRNA enrichment. **(B)** Volcano plot of gene enrichment in the “high” versus “low” reporter-expressing sorted fractions after infection with the H1 CRISPRi library (Table S2). Fold enrichment is calculated as the log_2_ of the mean counts for 3/5 sgRNAs targeting a given gene, while *p* values are calculated based on the distribution of all 5/5 sgRNA targeting a gene relative to the NTCs in the library. PKA subunit-encoding genes are shown in blue, GRKs are shown in red. **(C)** Enrichment of *ADRB2*-targeting sgRNAs in the each sorted population are plotted and color-coded based on the log_2_ enrichment of sgRNA counts as indicated in the legend. *ADRB2*-specific sgRNAs are colored in red, negative control sgRNAs are colored grey. sgRNAs used for validation experiments in Figure S2 are indicated. *p*-values were calculated by Mann-Whitney test using sgRNA activity relative to the negative control distribution.

### Genome-scale CRISPR interference screening identifies novel determinants of GPCR-mediated transcriptional signaling

We next carried out a genome-wide CRISPRi screen, testing the remaining libraries H2-H7 in CRE-GFP/CRISPR cells. Similar to H1, H2-H7 contain 5 sgRNAs/target gene as well as library-specific NTCs that are used for data normalization. As described above, CRE-GFP/CRISPR cells transduced with sgRNAs for 1 week were treated with isoproterenol for 4 h and sorted into three fractions. We detected sgRNAs for every gene targeted by our libraries (20,496 / 20,496 targeted transcription start sites corresponding to 18,903 unique genes), suggesting that there were no major bottlenecking effects due to counter-selection of genes with essential functions (Fig. 3a). Taking advantage of the large number of non-targeting control sgRNAs in our libraries, we generated “quasi-genes” by random sampling from non-targeting sgRNAs to empirically estimate a false discovery rate (FDR) for a given cutoff of gene “hit strength”, defined as the product of phenotype score and −log_10_(*p* value). We defined genes passing an FDR < 0.1 as hit genes. “Negative regulators” were identified as genes whose sgRNAs were enriched in the higher relative to lower fluorescence fractions, and “positive regulators” were identified as genes whose sgRNAs were enriched in the lower relative to the higher fluorescence fractions. Finally, pairwise comparison of sgRNA enrichment across the fractions allowed us to identify mediators with strong (genes which sgRNAs enriched in “high” vs “low” fraction or vice versa, Fig. 3a, Table S2) and weaker (genes which sgRNAs enriched in “medium” vs either “high” or “low” or vice versa, Fig. S3a) phenotypes with respect to cAMP signaling.

**Figure 3.**
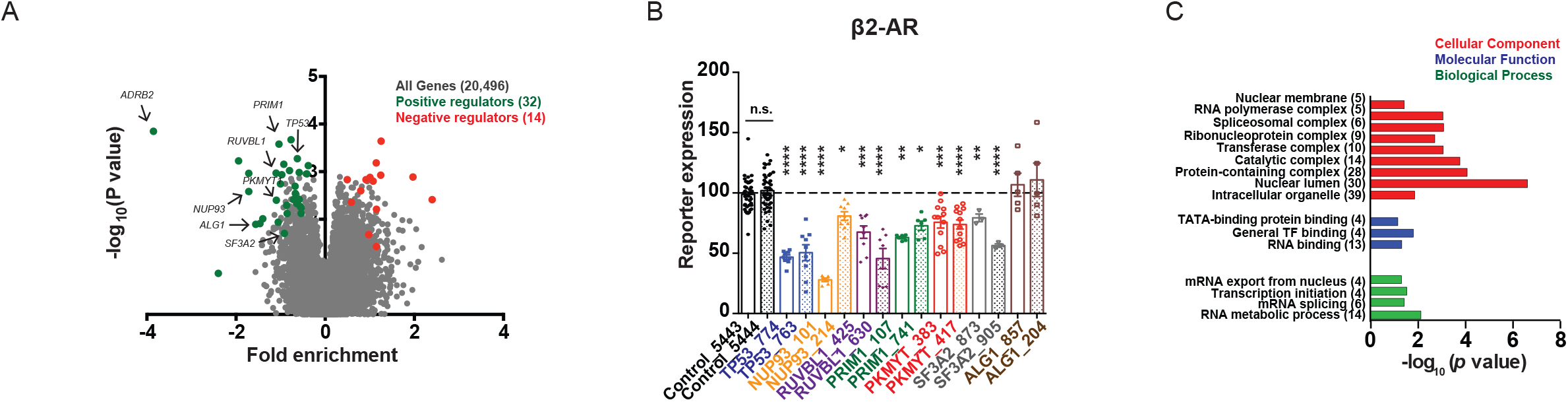
A high-throughput CRISPR-based genomic screen identifies novel modulators of GPCR signaling. **(A)** Volcano plot of gene enrichment in the “high” versus “low” fractions from all seven CRISPRi libraries. Significant hits with “hit strength” FDR ≤ 10% are color-coded based on their phenotype, and genes selected for follow-up validation are indicated with arrows. **(B)** Validation of the functional effects on β2-AR-dependent CRE-DD-GFP reporter upregulation for a subset of novel regulators identified by the CRISPRi screen. Two sgRNAs per gene were selected based on screen phenotype, individually cloned and tested in batches. Cells were treated with 1 μM Shield-1 and either 1 μM isoproterenol for 4 h and analyzed by flow cytometry. Mean GFP signal of the two control sgRNAs (5443 and 5444) for each batch was set to 100% and used to normalize the mean GFP values for all sgRNAs from that batch, as described in the “Materials and Methods” section. Data shown are mean from n = 3-12 per gene-specific sgRNA, and n = 41-42 for NTCs. **(B)** Enriched Gene Ontology categories (Table S5) among High-confidence hits identified through “high”/”low” comparison. Error bars = ± s.e.m. **** = *p* ≤ 0.0001; *** = *p* ≤ 0.001; ** = *p* ≤ 0.01; * = *p* ≤ 0.05 by one-way ANOVA test.

Using these criteria in pairwise comparisons across all sorted populations, we identified 46 hits from “high” vs “low” and 74 hits combined from “medium” vs “low” or “medium” vs “high” comparisons, 24 of which were shared. Next, we examined how likely are these hits to represent real regulators of receptor signaling. To evaluate the reliability of the screen results, we picked seven of the 46 genes with strong phenotypes (*ALG1, NUP93, PKMYT1, PRIM1, RUVBL1, SF3A2* and *TP53*) and two of the 50 unique genes with weaker phenotypes (*HRAS* and *MICAL3*) from four different CRISPRi sub-genome libraries annotated to carry out diverse cellular functions (Fig. 3a, S3a, Table 1), and examined the effects of depleting these genes on CRE-GFP transcription. In order to avoid possible clonal artifacts that may have disproportionally skewed the outcome in the original CRE-GFP/CRISPR cell line, we carried out the validation experiments in dCas9-KRAB-expressing cells that were co-infected with CRE-GFP and individual sgRNAs, and assayed the signaling responses in a polyclonal reporter-expressing cell population within a week. Following established protocols for validation of primary screens, we targeted each gene with two distinct sgRNAs.^20–22^ We found that depletion of 6/7 (86 %) of the hits identified from the “high”/”low” and 1/2 (50 %) of the hits from the “medium”/”low” gave rise to phenotypes consistent with the pooled screen result (Fig. 3b, S3b-c). Specifically, knockdown of the strong phenotype hits *NUP93, PKMYT1, PRIM1, RUVBL1, SF3A2* and *TP53* with two sgRNAs per gene significantly blunted the isoproterenol-dependent CRE-GFP induction seen in control cells (Fig. 3b), while silencing of *HRAS* led to a reproducible increase in reporter transcription across the sgRNAs tested, which was statistically significant for sgRNA_187 (Fig. S3b). Because only 1 out of the 2 genes tested from the “medium”/”low” comparison was consistent between the follow-up analysis and the pooled screen, we designated this set of genes as “Putative hits” (Table S3), and caution that this set of hits has a higher probability of false positives. In contrast, the high true positive rate amongst genes identified from “high”/”low” sorted fraction comparisons suggests that a significant fraction will be bona fide novel regulators of GPCR/cAMP signaling, and we defined these as “High-confidence hits” (Table S4).

**Table 1.** Hits from the screen selected for validation analyses.

| **Gene** | **Localization<sup>16</sup> and Function<sup>43</sup>** | **Role(s) in gene expression regulation** |
| --- | --- | --- |
| *HRAS* * | Plasma membrane and cytosol; GTPase, activation of signal transduction | Via Ras cascade activation (c-Raf, PI3K) |
| *NUP93* * | Nucleus; Nuclear pore assembly and maintenance | Smad4 nuclear translocation<sup>27</sup>; binds to active chromatin<sup>26,28,29</sup> |
| *PKMYT1* * | ER, Golgi; Tyrosine- and threonine-specific kinase; Golgi fragmentation, cell cycle regulation | Regulates β-catenin/Wnt signaling by phosphorylating and inactivating GSK3beta<sup>31</sup> |
| *PRIM1* * | Vesicles and nucleus; DNA primase | Involved in DNA replication |
| *RUVBL1* * | Nucleus; DNA-dependent ATPase and helicase | Chromatin remodeling; can drive oncogenesis as a co-factor for Myc and β-catenin<sup>44,45</sup> |
| *SF3A2* * | Nucleus; Pre-mRNA splicing factor | Component of the spliceosome |
| *TP53* * | Nucleus; Tumor suppressor p53 | Transcription factor; forms complex with CBP/CREB<sup>46,47</sup> |
| *ALG1* | ER and nucleoli; biosynthesis of lipid-linked oligosaccharides | No known direct role(s) |
| *MICAL3* | Plasma membrane, cytosol and nucleus; actin filament disassembly, vesicle trafficking | No known direct role(s) |

\* = validated in follow-up analysis

Depletion of *ADRB2* yielded the most extreme effect on GPCR-dependent induction of the reporter across all libraries tested-an expected result validating our experimental approach (Fig. 3a). In addition to the receptor itself, three other effectors are well known to be involved in this pathway. Gαs (encoded by *GNAS*) and CREB (encoded by *CREB1*) are required for signal initiation and transcriptional activation of target genes in the nucleus, respectively, and thus would be predicted to blunt CRE-GFP accumulation when depleted. Yet, neither of these mediators scored as a hit in the screen. Since pharmacological inhibition of CREB blunted the forskolin-induced accumulation of the reporter in our earlier experiments (Fig. 1c), we reasoned that the most likely explanation for this unexpected phenotype is low activity of the sgRNAs targeting these genes. Indeed, retests of individual *CREB* and *GNAS* sgRNAs confirmed that the resulting cell lines retained robust CRE-GFP up-regulation upon β2-AR stimulation (Fig. S4a-b), and quantitative PCR analysis showed very minimal *CREB* and *GNAS* mRNA depletion likely insufficient to interfere with protein function (Fig. S4c). Beta-arrestin 2 (encoded by *ARRB2*), another recognized GPCR regulator, is a cytosolic adaptor canonically known for its role in shutting off G protein-dependent signaling by uncoupling the GPCR-G protein complex at the plasma membrane and facilitating subsequent receptor internalization.^14^ While previous work has shown that *ARRB2* depletion leads to more prolonged cAMP production due to receptors that fail to desensitize^14^, the role of arrestin in GPCR-induced transcriptional responses has not been explored. Arrestin was not identified as a hit in the genomic screen. Follow-up experiments using an individually cloned sgRNA corroborated that *ARRB2* is necessary for GPCR desensitization and endocytosis (Fig. S4c-e) but is dispensable from cAMP responses generated through direct cyclase stimulation with forskolin (Fig. S4f). Yet, arrestin depletion had no effect on CRE-GFP accumulation (Fig. S4b). This lack of phenotype did not reflect an artifact of the reporter assay, because *ARRB2* knockdown had no effect on transcriptional upregulation of two known endogenous CREB target genes, *PCK1* and *NR4A1*, either (Fig. S4g).^7^ These data therefore support the results from the screen and suggest that knockdown of arrestin does not impact activation of the β2-AR-dependent transcriptional responses.

### Factors identified through the screen regulate distinct steps of the GPCR/cAMP pathway

We next focused on the high-confidence hits identified through the screen, which comprised 32 positive and 14 negative regulators (Table S4). Interestingly, with the exception of *ADRB2*, none of the remaining 45 genes have been previously shown to mediate aspects of GPCR/cAMP signaling. Functional classification using the Gene Ontology Resource^23–25^ revealed a disproportionate enrichment for genes encoding regulators of transcription and translation (Fig. 3c, Table S5). Because the transcriptional reporter is compatible with flow cytometry, it allowed us to examine the impact of the novel factors on multiple cell states in parallel in order to dissect their specificity and mechanisms of action. First, by assaying CRE-GFP expression in uninduced cells we evaluated whether the validated hits regulate the pathway only upon activation of β2-AR/cAMP signaling, or if they also play roles under basal conditions. We observed gene-specific effects on basal transcription. Knockdown of the majority of genes did not reproducibly impact reporter levels in resting cells across the two gene-specific sgRNAs tested (Fig. S5a). We found two exceptions-*PKMYT1* and *SF3A2. PKMYT1* depletion led to a mild (~1.3-fold) but statistically significant increase in basal reporter levels with both sgRNAs (Fig. S5a). Knockdown of *SF3A2* resulted in a much more dramatic upregulation of basal CRE-GFP (~2.5-7-fold increase relative to controls, highly significant for one of the two sgRNAs, *p* < 1.0×10^-3^ by one-way ANOVA test) (Fig. S5a). This phenotype was unexpected given the *blunted* isoproterenol-dependent response in the same cells (Fig. 3b). While this intriguing result suggests differential involvement of *SF3A2* in mediating resting versus signaling states, we noticed that knockdown with one of the two sgRNAs (sgRNA_873) led to increased cell death. Although the other sgRNA (sgRNA_905) did not have discernible effects on cell fitness but yielded the more striking phenotype, our data cannot rule out the possibility that the observed changes in cAMP signaling reflect pleiotropic effects on cell viability.

Given the factor-specific effects on induced and basal reporter expression, we next asked if the modulators are selective for the β2-AR cascade by stimulating cAMP production through endogenous adenosine receptors (A2-Rs) instead. With all six genes, we saw significantly diminished transcription upon cAMP activation in cells stimulated with the agonist NECA that mirrored the isoproterenol-dependent responses, indicating that many of the modulators identified by the screen are likely to be of universal significance to the cAMP cascade (Fig. S5b). To dissect whether these factors act at the GPCR or are involved in downstream step(s) of the pathway, we next assessed CRE-GFP induction when AC activity was stimulated directly with forskolin. We found that depletion of five genes (*TP53, NUP93, PRIM1, RUVBL1*, and *SF3A2*) also significantly blunted forskolin-induced reporter accumulation with at least one of two sgRNAs tested. Interestingly, *PKMYT1* knockdown in forskolin-stimulated cells led to very mild decrease in CRE-GFP expression only for one of the sgRNAs, while the other sgRNA had no effect on the reporter (Fig. S5c). In contrast, both *PKMYT1* sgRNAs diminished the transcriptional responses to β2-AR and A2-R activation (Fig. 3, S5b), suggesting that this kinase may disproportionately impact GPCR-dependent signaling.

Based on the phenotypes that the novel factors displayed in the validation reporter assay, we picked *NUP93*, which had a robust effect on transcription across different activation conditions, and *PKMYT1*, which preferentially affected the GPCR-induced responses, for more comprehensive mechanistic characterization. *NUP93* encodes a scaffold component of the nuclear pore complex that regulates gene expression both directly, by facilitating nucleocytoplasmic transport of signaling molecules, and indirectly through association with select chromatin regions.^26–29^ The *PKMYT1* gene codes for a ER/Golgi-associated serine/threonine kinase that was first identified as a cell cycle regulator (via phosphorylation and inactivation of Cdc2)^30^, but more recently was shown to have additional substrates. Notably, Pkmyt1 was found to phosphorylate GSK3beta and, accordingly, to mediate beta-catenin signaling.^31^ We first verified the efficiency of sgRNA-mediated knockdowns and their effects on endogenous transcriptional signaling with quantitative PCR. While both *PKMYT1* and *NUP93* silencing (Fig. S6a) impacted isoproterenol-induced upregulation of endogenous CREB targets, only *NUP93* was also required for the forskolin responses (Fig. 4a and S6b), confirming the reporter result. We next interrogated the involvement of these factors in cAMP production. To evaluate second messenger production in induced cells, we took advantage of a luciferase-based cAMP biosensor that enables second messenger detection in real time. Mirroring the CRE-GFP reporter phenotype, we saw a significant reduction in total cAMP produced in *NUP93* knockdown cells upon AC stimulation through the β2-AR, the A2-R or directly with forskolin (Fig. 4b, right). Interestingly, we observed that *NUP93* depletion impacts not only induced but also basal cAMP levels measured with a high-sensitivity colorimetric immunoassay (Fig. 4b, left and S6c). These results suggest that the diminished transcriptional responses that we observed in *NUP93*-deficient cells are a consequence of reduction in upstream cAMP accumulation. In contrast, *PKMYT1* knockdown had no impact on bulk cytosolic cAMP accumulation (Fig. S6d).

**Figure 4.**
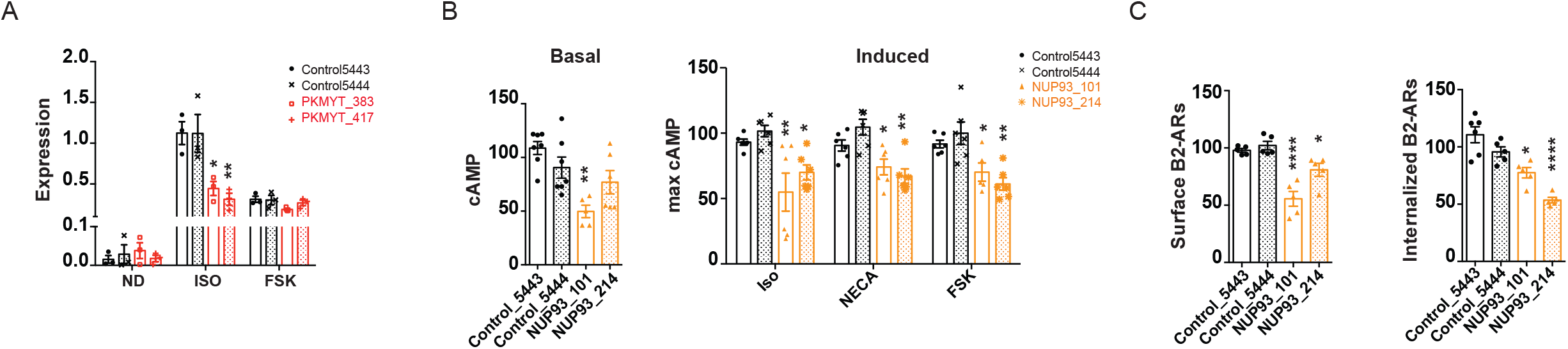
Novel hits identified by the screen regulate GPCR/cAMP signaling through distinct mechanisms. **(A)** *PKMYT1* depletion blunts isoproterenol-dependent accumulation of an endogenous CREB target gene, *PCK1*. Cells were left untreated (basal, ND = no drug), treated with 1 μM isoproterenol, or with 10 μM forskolin 1 h, RNA was isolated, reverse transcribed and analyzed by qPCR. Data plotted are *GAPDH*-normalized mean levels from n = 3. **(B)** *NUP93* knockdown diminished cAMP production following GPCR-dependent and direct induction of adenylyl cyclase enzymes. Basal cAMP levels (left) were measured by cAMP ELISA assay, and induced levels (right) were measured using a luciferase-based biosensor in real time upon addition of 1 μM isoproterenol, 10 μM NECA or 10 μM forskolin (FSK). ELISA values were normalized to total protein per sample, and maximum biosensor values were normalized to account for plasmid transfection efficiency (see “Materials and Methods”). All values shown are percent of averaged controls. Data are mean from n = 4-5 (ELISA) and n = 6 (biosensor). **(C)** Effects of *NUP93* depletion on trafficking of β2-AR analyzed by flow cytometry. Cells transiently transfected with flag-tagged β2-AR were either left untreated or were induced with 1 μM isoproterenol for 20 min, and cell-surface levels of flag-tagged receptor were measured by incubation with Alexa-647-M1 on ice and flow cytometry analysis. Average control values were set to 100% and used to normalize the data as described in the “Materials and Methods” section. Data are mean from n = 5. Error bars = ± s.e.m. **** = *p* ≤ 0.0001, ** = *p* ≤ 0.01, * = *p* ≤ 0.05 by one-way ANOVA test.

One highly regulated cellular process that can impact the signaling outputs of the pathway is GPCR trafficking. The receptor itself has to be delivered to the plasma membrane upon translation and subsequently traffics through the endocytic/recycling pathway during activation.^32^ Therefore, we next focused on the roles of Nup93 and Pkmyt1 in β2-AR trafficking to and from the plasma membrane. Using flow cytometry to assay cell surface levels and drug-induced internalization rates of flag-tagged β2-ARs transiently expressed in CRISPRi cells, we observed a *NUP93*-dependent decrease in both (Fig. 4c). Notably, the diminished β2-AR cell surface expression was not due to downregulation at the mRNA level (Fig. S6e), and the effect was *NUP93*-specific, as knockdown of *PKMYT1* did not affect β2-AR cell surface expression or internalization (Fig. S6f). Thus, nucleoporin function is required both for proper export of the GPCR to the plasma membrane and for ligand-induced internalization. This aberrant localization of the receptor in *NUP93*-depleted cells may explain the diminished basal and induced cAMP production (Fig. 4b and S6c, see “Discussion”). Meanwhile, *PKMYT1* was not required for cAMP production or receptor trafficking, which suggests that the kinase acts at a downstream step to regulate CREB activity in a GPCR-dependent manner.

## Discussion

Here, we present an unbiased CRISPR-based screening platform that can be used to identify both positive and negative regulators of GPCR/cAMP signaling. To our knowledge, this is the first functional genomic screen of a GPCR or second messenger cascade. We circumvented the lack of robust readouts that has obstructed the application of such approaches to these pathways by generating and optimizing a fluorescent transcriptional reporter that can be coupled with sorting-based readouts (Fig. 1 and 2). While the CRISPR approach described here has certain limitations arising from incomplete gene silencing or compensatory mechanisms allowing the cell to maintain signaling (Fig. 2b, S4), it is a valuable tool to dissect these complex signaling pathways in an unbiased and systematic way.

Due to the sortable nature of the platform, we were able to assay regulators with strong and weaker phenotypes by analyzing sgRNA distribution among three fractions with distinct reporter expression (“high”, “medium” and “low”). Based on experimental validation rates, we have designated genes identified through pairwise comparisons involving the “medium” sorted population of cells as “Putative hits” (Table S3) and genes identified in the “high” vs “low” comparison as “High-confidence hits” (Table S4). Specifically focusing on genes with weaker phenotypes (50 unique hits), in follow up experiments we show that depletion of *HRAS* yielded an effect on CRE-GFP expression, while knockdown of *MICAL3* did not (Fig. S3). Therefore, while genes in this set represent a valuable list of potential mediators to inform future studies of the pathways based on the validation of *HRAS* and the overall extensive overlap between genes with strong and weaker phenotypes (24 shared hits, *p* < 1 x 10^-52^ by Fisher’s exact test), they have a higher false positive likelihood. In contrast, genes with strong phenotypes in the screen (46 hits) are very likely to present bona fide regulators, as we have successfully validated six of them on a gene-by-gene basis (86 %, Fig. 3b, Table S4). Notably, with the exception of the β2-AR-encoding gene, *ADRB2*, the remaining 45 “High-confidence hits” represent factors not previously known to mediate GPCR or cAMP signaling.

Because the experimental platform scores transcriptional activation, a downstream output of the GPCR/cAMP cascade, it is poised to identify functional regulators across multiple steps of the pathway. Consistent with that, novel hits are annotated to localize to different compartments (Fig. 3c), and we have shown that they impact signaling through distinct mechanisms. Some genes are required for transcription under both basal and induced conditions (*SF3A2*), while others play a regulatory role only upon activation of the cascade (*PKMYT1, TP53, RUVBL1, NUP93* and *PRIM1*) (Fig. 3b, S5). In addition, we found a factor, *PKMYT1*, which disproportionally impacted transcriptional responses activated via GPCRs but not by direct AC stimulation (Fig. 3b, 4a, S5b-c). Since one obvious point of divergence between pathways activated through a receptor and with forskolin would be upstream of or at the level of the cyclase itself, we anticipated that Pkmyt1 might regulate these early steps. Interestingly, our data argue that this is not the case, as neither β2-AR trafficking nor bulk cytosolic cAMP production was affected by *PKMYT1* knockdown (Fig. S6d,f). This suggests that the kinase impacts a more downstream stage(s). Pkmyt1 is associated with the Golgi^30^, where the Type II PKA regulatory isoform is also predominantly localized.^33^ Thus, it is possible that the kinase regulates PKA activity and/or subsequent catalytic subunit release/nuclear translocation in a receptor-dependent manner (Fig. 5). Alternatively, signaling from endosomal receptors has emerged as a new paradigm in the field. Specifically, it has been demonstrated that GPCR/cAMP-dependent transcriptional programs are initiated selectively from endosomal receptors.^7,34^ Since we see a pathway-selective kinase effect specifically on transcription, another tentative mechanism could involve Pkmyt1-dependent modulation of endosomal effector localization and/or function. Future proteomics studies focused on elucidating the phosphotargets of Pkmyt1 would be instrumental in shedding light on the precise mechanisms.

**Figure 5.**
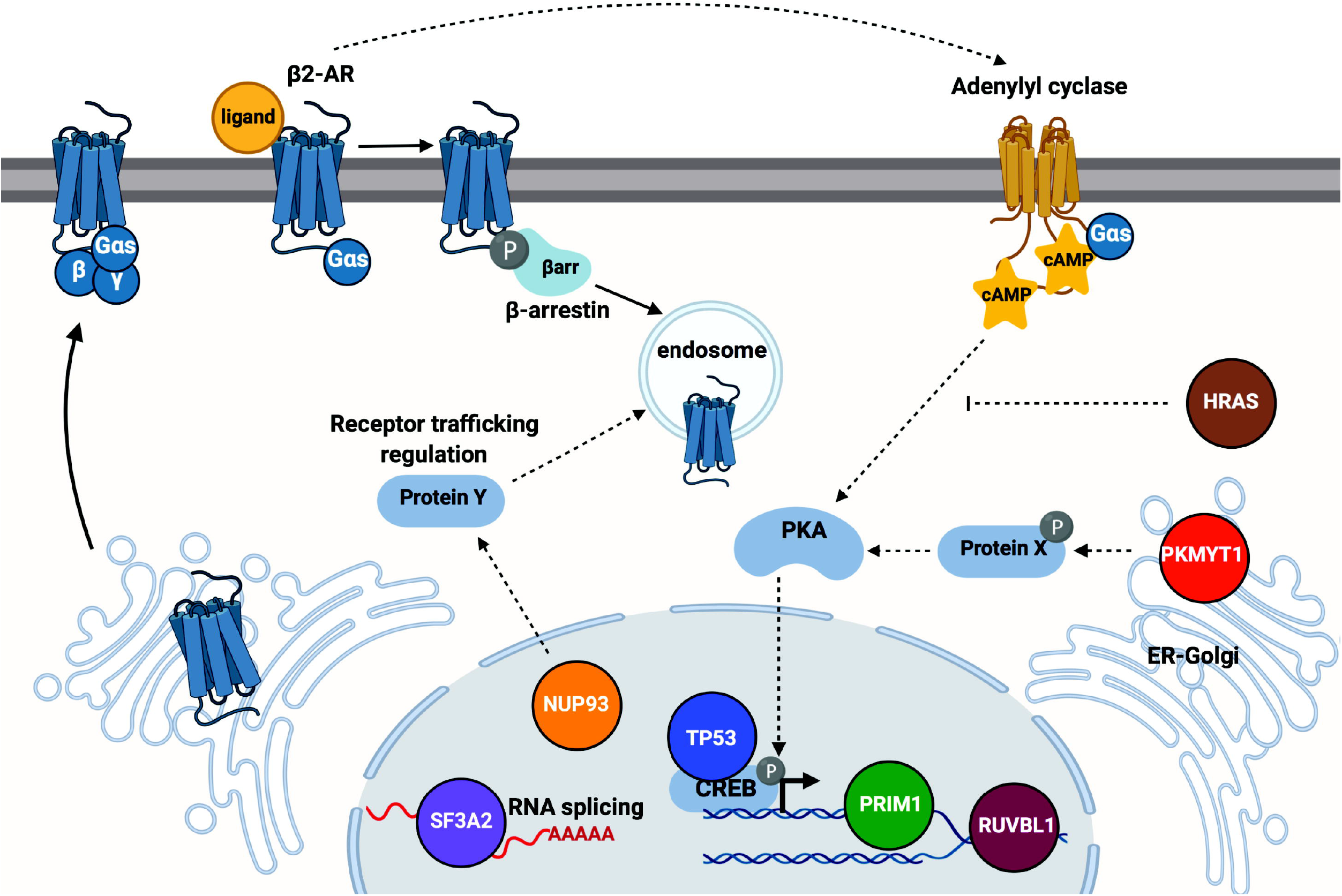
Model depicting proposed regulatory roles for validated GPCR/cAMP mediators. Proposed roles are assigned based on functional and localization annotations of the factors (Table 1) and experimental evidence provided by this study.

In contrast to Pkmyt1, our analyses revealed that another factor, the nucleoporin-coding gene *NUP93*, impacts a number of steps in the signaling cascade, including receptor trafficking to and from the plasma membrane, cAMP production, and transcriptional responses (Fig. 4). Because *NUP93* depletion affected not only induced but also basal cAMP, we propose that the primary effects of the nucleoporin are on GPCR trafficking, and that the aberrant localization of the receptor consequently underlies the blunted cAMP production and transcriptional responses (Fig. 5). Furthermore, *NUP93* was necessary both for receptor-dependent and independent cAMP accumulation and transcription (Fig. 3–4 and S5-6). Thus, it is very likely that this factor impacts the intracellular localization of additional signaling components, including adenylyl cyclase enzymes. In agreement with this model, GPCRs and transmembrane ACs require some of the same machinery to traffic through the endocytic pathway and are frequently found in numerous intracellular compartments together.^35–39^ Moreover, a recent study reported that a cyclase co-internalized with the β2-AR upon its activation, which further underscores the shared regulatory mechanisms between these signaling effectors.^38^ Based on this, we propose that Nup93 regulates the expression and/or function of endocytic trafficking components critical for proper GPCR/cAMP localization and signaling (Fig. 5).

A couple of known mediators of receptor function did not yield significant phenotypes in the screen. We showed that for some genes (CREB, Gαs) this is due to low sgRNA activity (Fig. S4c). For others (PKA, GRKs, adenylyl cyclase enzymes), functional effects can be masked by genetic redundancy as the current screen relies on silencing of individual genes. Indeed, we observed a modest enrichment of sgRNAs for two of the PKA subunits (RIb and RIIa) and one of the GRKs (GRK2), but not the others (Fig. 2b). These results are consistent with phenotypes reported by other studies utilizing single gene knockdowns of these effectors.^18,19,38^ A strategy to overcome such redundancies and to enable fine-mapping of new mediators to distinct steps within the pathway would be to utilize combinatorial gene silencing or to test the effects of simultaneous gene depletion/gene activation on GPCR-dependent signaling. In fact, CRISPRi has been used successfully both for creating loss-of-function genetic interaction maps^40^ and in conjunction with CRISPR activation (CRISPRa)^15^ in the past, and thus the screening platform described here should be easily amenable to either experimental set up for future analyses. Still, other known GPCR regulators did not score in the assay for biological reasons. While it is well recognized that beta-arrestin is required to turn off G protein-dependent signaling and to facilitate GPCR internalization^14^, to our knowledge it is not established how the adaptor impacts GPCR/cAMP-dependent nuclear responses. Consistent with previous studies, we observed that CRISPRi depletion of arrestin with an individually cloned sgRNA resulted in more persistent cAMP signaling and defective GPCR endocytosis (Fig. S4d-e). However, lack of arrestin did not impact the transcriptional responses as evidenced in a CRE-GFP reporter assay and by quantitative PCR analysis of endogenous CREB targets (Fig. S4b,g). An intriguing possibility to account for the divergent upstream and downstream effects of arrestin (and GRKs) would be that the cell has evolved mechanisms to “buffer” minor fluctuations in accumulated cAMP levels. Indeed, in a recent study of the β2-AR pathway we reported that the CREB-dependent transcriptional responses are uniform across a gradient of cAMP concentrations.^41^ In addition, the screen described here was carried out by stimulation of the receptor with a single dose of isoproterenol over the course of 4 hours. The main regulators of GPCR desensitization, arrestin and GRKs, have more pronounced effects on cAMP kinetics upon repeated receptor stimulation and over a shorter timecourse of activation.^13,14^ We anticipate that altering the induction parameters used in the assay may change the magnitude of the effects and can thus be used as an additional experimental strategy to uncover new layers of regulation of these pathways.

A significant fraction of the novel hits identified in this study, including five of the validated genes, encode either nuclear and/or components of known DNA- and RNA-binding complexes (Fig. 3c, Table 1). Likely, some of these may regulate the pathway indirectly, by controlling the expression of other more proximal effectors of GPCR/cAMP signaling. For example, among the hits encoding RNA-binding proteins, we found a highly enriched cluster of splicing factors and splicing factor-interacting proteins (Fig. 3c). We confirmed that depletion of one of the novel hits, *SF3A2*, encoding a subunit of the SF3A splicing complex of the mature U2 snRNP, led to *blunted* transcriptional response upon cascade activation, but a paradoxical *increase* in basal transcription (Fig. 3b, S5). This result is striking given that both the β2-AR and the transcriptional reporter are intronless and, thus, not direct subjects to splicing. If not the receptor, an attractive set of candidates for spliceosomal targets would be phosphodiesterases (PDEs), which hydrolyze cyclic nucleotides and are thus critical regulators of cellular cAMP levels (Fig. 5). PDE function is extensively regulated by splicing, allowing the 21 human PDE genes to express more than 200 distinct protein isoforms.^42^ This study provides a pivotal starting point to begin to dissect the underlying regulatory logic, but future work will delineate the specific targets of *SF3A2* and of the other nuclear factors identified in the screen, and identify the effectors that are directly involved in the control of basal and induced states of the cAMP cascade.

In summary, we have developed a high-throughput screening paradigm for unbiased systematic dissection of GPCR/cAMP cascades, and provide proof of principle that this platform can be applied to successfully identify bona fide regulators of β2-AR signaling. We envision that this reporter-based platform will aid the high-throughput dissection of other important aspect of GPCR signal transduction in the future, including identification of regulators of drug-specific responses, functional characterization of receptor features, and identification of novel chemical compounds that modulate receptor activity. Thus, it can become an invaluable tool to illuminate drug-dependent mechanisms and pinpoint candidates for pharmacological and genetic manipulation of these essential pathways.

## Acknowledgments

We thank members of the Tsvetanova, von Zastrow and Kampmann laboratories for discussions and critical feedback on the manuscript. Grace Peng (von Zastrow laboratory, UCSF) cloned and kindly provided the pGLO-20F/Rluc vector for real-time analysis of cAMP levels. Figure 5 was created with BioRender.com. This work was supported by the National Institute of Health [Grants MH109633 to N.G.T, DA010711 and DA012864 to M.v.Z., and DP2 GM119139 to M.K.]

## Author Contributions

N.G.T designed the experiments. K.S. and N.G.T. carried out the experiments. R.T. prepared the CRISPR libraries and designed the computational pipeline for CRISPRi screen analysis. K.S. and N.G.T. wrote the manuscript. K.S., R.T., M.v.Z., M.K. and N.G.T. edited the manuscript.

## Materials and Methods

### Chemicals

(-)-Isoproterenol hydrochloride was purchased from Sigma-Aldrich (Cat #I6504), dissolved in water/100 mM ascorbic acid to 10 mM stock, and used at 1 μM final concentration. 5’-(N-Ethylcarboxamido)adenosine (NECA) was purchased from Sigma-Aldrich (Cat #119140), solubilized in DMSO to 10 mM stock, and used at 10 μM final concentration. Forskolin was purchased from Sigma-Aldrich (Cat #F6886), dissolved in DMSO to 10 mM stock, and used at 10 μM final concentration. Shield-1 ligand for stabilization of DD-tagged proteins was purchased from Takara Bio (Cat # 632189) and added to the cell medium to 1 μM final concentration. The cell-permeable cAMP analog, 8-Bromo-cAMP, was purchased from Santa Cruz Biotechnology (Cat #sc-201564), dissolved in DMSO and used at 1 mM final concentration within 1 month of resuspension. The CREB inhibitor compound, 666-15, was purchased from Tocris Bioscience (Cat #5661), resuspended in DMSO and used at 100 nM final. At doses higher than 100 nM, 666-15 was toxic in our cell line. H89 dihydrochloride hydrate was purchased from Sigma-Aldrich (Cat #B1427), resuspended in DMSO to 10 mM stock, and used at 10 μM final concentration. D-luciferin sodium salt (Cat #LUCNA) and coelenterazine (Cat #CZ) were purchased from GoldBio and resuspended to 100 mM in 10 mM Hepes, and 10 mM in ethanol, respectively, and stored protected from light.

### Construct cloning

A lentiviral plasmid encoding a transcriptional reporter for CREB activity was generated by PCR amplification of a 2xCRE promoter-driven GFP N-terminally tagged with the ProteoTuner destabilization domain (CRE-DD-GFP, Takara Bio, Cat #631085) and Gibson cloning into the FUGW lenti-vector backbone (Addgene, Cat #14883) to replace the pUbc-GFP fragment. The vector expressing a catalytically dead Cas9, dCas9, fused to BFP and a KRAB repressor domain (pHR-SFFV-dCas9-BFP-KRAB) and the parental vector for sgRNA expression under a U6 promoter (pU6-sgRNA-EF1alpha-puro-T2A-BFP) were a gift from Jonathan Weissman (Addgene, Cat# 46911 and #60955, respectively) and were described previously.^15,48^ sgRNAs were cloned by annealing complementary oligonucleotides purchased from IDT (Table S6) and ligation into a BstXI/BlpI digested pU6-sgRNA-EF1alpha-puro-T2A-BFP backbone as described previously.^15^ The cAMP luminescence biosensor with renilla luciferase, pSF-CMV-GloSensor20F-IRES-Rluc (pGLO-20F/Rluc), was generated by Gibson cloning using EcoRI/EcoRV-digested backbone from pSF-CMV-EMCV-Rluc (Boca Scientific, Cat #OG296) and the sequence encoding a genetically modified firefly luciferase into which a cAMP-binding domain has been inserted from the pGloSensor-20F (Promega, Cat #E1171).

### Cell culture and reporter cell line generation

HEK293 and HEK293T cells were obtained from ATCC and grown at 37°C/5% CO_2_ in Dulbecco’s Modified Eagle Medium (4.5 g/L glucose and L-glutamine, no sodium pyruvate) from Thermo Fisher Scientific (Cat #11965118) supplemented with 10% fetal bovine serum. Clonal HEK293 cells stably expressing pCRE-DD-zsGreen1 and dCas9-BFP-KRAB were generated by lentiviral transduction and fluorescence activated cell sorting for GFP and BFP on a BD FACS Aria 2 (BD Biosciences). Individual clones were expanded and tested for 1) high cAMP-induced GFP expression, and 2) dCas9 transcriptional repression with sgRNAs against the *TRFC* and *ADRB2* genes. For these experiments, individual clones HEK293 cells stably expressing the cAMP reporter and dCas9 were seeded on 6-well dishes at ~30% confluence. The next day, the cells were transduced with gene-specific sgRNAs or non-targeting control sgRNAs. For *TFRC* knockdown experiments, cells were expanded for 4 days after lentivirus transduction. To quantify knockdown efficiency, ~1 x 10^5^ cells were lifted in PBS/EDTA (Fisher Scientific, Cat #BWBE02017F), washed once in PBS/5% FBS, resuspended in 100 μl PBS/5% FBS and blocked with 2.5 μg of Fc-blocking solution (BD Bioscience, Cat #564220) for 15 min at room temperature. Cells were then stained for transferrin receptor expression by adding 0.25 μg of PE/Cy7 antihuman CD71 [CY1G4] (Biolegend, Cat# 334111) directly to the blocking solution for 30 min at room temperature. Excess antibody was removed by two washes in PBS/5% FBS, cells were re-suspended in 150 μl of PBS/5% FBS and analyzed on a BD FACS Canto2 flow cytometer (BD Biosciences) for PE-Cy7 expression. For *ADRB2* knockdown experiments, 48 h after transduction with the sgRNAs, cells were selected with 1 μg/ml puromycin for 2 days, then recovered and expanded for 3 days in regular medium without antibiotic. To test loss of receptor function, cAMP signaling was activated either with 1 μM isoproterenol or 1 mM 8-Br-cAMP in medium containing 1 μM Shield-1 for 4 h, and cells were resuspended in 100 μl PBS for flow cytometry analysis of GFP expression on a BD FACS Canto2.

### Lentivirus production

For production of lentivirus used in the genomic CRISPRi screen, HEK293T cells were transfected with sgRNA library lentivirus vectors and standard packaging vectors (VSVG and psPAX2) using *Trans*IT-LTI transfection reagent (Mirus, Cat# 2306) following previously published protocols.^17^ Supernatant was harvested 72 h after transfection and filtered through a 0.45 μm SFCA filter. The harvested virus was used immediately to transduce the CRE-GFP/CRISPR reporter cell line. For production of lentivirus used in individual sgRNA testing, HEK293T cells were transfected with sgRNA lentivirus vector and standard packaging vectors (VSVG and psPAX2) using Lipofectamine-2000 transfection reagent (Thermo Fisher, Cat# 11668027) following recommended protocols. Supernatant was harvested 72 h after transfection and filtered through a 0.45 μm SFCA filter. The harvested virus was either used on the same day or concentrated and snap-frozen before use.

### Pooled CRISPRi genomic screening

CRISPRi libraries used in this study were described previously.^49^ CRE-GFP/CRISPR cells were infected with sgRNA libraries (Table S1) at an effective MOI of < 1 sgRNA/cell. Two days after infection, puromycin was added to 0.85 μg/ml final concentration for 72 h to select infected cells. Then, cells were allowed to recover in medium without puromycin for 3 days. On the day of the experiment, isoproterenol and Shield-1 were added to 1 μM final concentration each for 4 h. Cells were washed in ice-cold PBS, lifted in ice-cold Trypsin/0.25% EDTA and resuspended in PBS/2% FBS for sorting on a BD FACS Aria2 based on fluorescent reporter expression (GFP), gating for BFP+ (sgRNA+) singlets. We divided the cells into three fractions (high, medium, low) each collected in a separate tube. Invariably upon β2-AR stimulation, we observed a large right-shifted GFP peak (corresponding to cells with active reporter accumulation) and a small left-shifted peak (corresponding to cells with less or no reporter transcription). We set the “low” fraction gate at the boundary between these two peaks and collected the left-shifted population of cells (“low”), and divided the right-shifted peak into two equal fractions which comprised the “medium” and “high” populations. The sorted cell:library elements ratio was maintained at >= 100 to ensure sufficient sgRNA coverage. Genomic DNA was isolated from FACS-sorted populations using QIAamp DNA Blood Maxi kit (Qiagen, Cat #51192), and PCR-amplified with Q5 Hotstart high-fidelity 2X mastermix (NEB, Cat #M0494) and barcoding primers (Table S7) for 27 cycles. PCR reactions were purified on QIAquick PCR purification columns (Qiagen, Cat #28106) and loaded onto a 20% polyacrylamide/TBE gels. A product band ~270 bp was excised and DNA was extracted, quantified and sequenced on an Illumina HiSeq-4000 using custom primers (Table S7).

### Bioinformatic analysis

Screens were analyzed as previously described and the analysis script is available for download from https://kampmannlab.ucsf.edu/volcano-plot.^17^ Briefly, raw sequencing reads from next-generation sequencing were cropped and aligned to the reference using Bowtie^50^ to determine sgRNA counts in each sample. The following analysis steps were carried out on a library-by-library basis. First, Log_2_ fold change (L_2_FC) was calculated for each sgRNA between two samples for comparison. Next, gene level knockdown phenotype scores (epsilon) were determined by averaging LFCs of the most extreme 3 sgRNAs targeting this gene. The statistical significance for each gene was determined by comparing the set of *p* values for sgRNAs targeting it with the set of *p* values for non-targeting control sgRNAs using the Mann-Whitney U test. To correct for multiple hypothesis testing, we also generated ‘negative-control-quasi-genes’ by random sampling of 5 with replacement from non-targeting control sgRNAs and calculated knockdown phenotype scores and *p* values for each of them as described above. Then, the “hit strength”, defined as the product of knockdown phenotype score and −log (*p* value), was calculated for all genes and ‘negative-control-quasi-genes’. We calculated the false-discovery rate (FDR) based on the distribution of all hit strength scores, and a cutoff was chosen based on an FDR < 0.10. Enriched functional categories among High-confidence target genes shown in Fig. 3c were identified using the Gene Ontology Resource (geneontology.org)^23–25^ using the Benjamini-Hochberg False Discovery Rate (FDR < 0.05).

### Individual retests of select sgRNAs

For all follow-up experiments, 2 control sgRNAs (5443 and 5444) and 2 sgRNAs targeting each gene of interest (Table S6) we cloned into the parental vector for sgRNA expression under a U6 promoter (pU6-sgRNA-EF1alpha-puro-T2A-BFP) at the BlpI/BstXI sites. The gene-specific sgRNAs were tested in batches and the same set of 2 control sgRNAs was included in each batch for normalization purposes. HEK293 stably expressing dCas9-BFP-KRAB were seeded on 6-well dishes at ~30% confluence and transduced with viral supernatant containing the sgRNAs of interest and pCRE-DD-zsGreen1, respectively. Two days post infection, cells were selected with 1 μg/mL puromycin for 48 h, then recovered in DMEM + 10% FBS without antibiotic for three days prior to assaying. For all experiments, statistical significance was calculated between gene-specific sgRNAs and matched NTCs within their experimental batches by one-way ANOVA test. Our criteria for reporting an sgRNA effect as significant requires that the *p* < 0.05 with respect to both NTC sgRNAs.

#### Flow cytometry-based experiments with pCRE-DD-zsGreen1

Cells were treated with 1 μM Shield-1 ligand alone (basal) or with 1 μM Shield-1 ligand and either 1 μM isoproterenol, 10 μM forskolin, or 10 μM NECA (induced) for 4 hours. We analyzed 5,000-10,000 total cells per sample using a BD FACS Canto2 instrument and gated for sgRNA-expressing (BFP+) singlets. From these measurements, the GFP mean of the control sgRNAs was averaged and set to 100% for each treatment condition, and the GFP mean for each sgRNA was normalized to this value (expressed as % of mean control value).

#### qPCR analysis of target gene expression

Cells were treated with 1 μM isoproterenol for 1 h, and total RNA was extracted from the samples using the Zymo Quick-RNA Miniprep Kit (Genesee, Cat #11-327) or Qiagen RNeasy Mini Kit (Qiagen Cat #74106). Reverse transcription was carried out using iScript RT supermix (Biorad, Cat #1708841) or Superscript II Reverse Transcriptase (ThermoFisher Scientific, Cat #180644022) following recommended manufacturer protocols. The resulting cDNA was used as input for quantitative PCR with CFX-384 Touch Real-Time PCR System (Biorad). Power SYBR Green PCR MasterMix (ThermoFisher Scientific, Cat #4367659) and the following primers were used for the qPCR reactions-*GAPDH*: F 5’-CAATGACCCCTTCATTGACC-3’ and R 5’-GACAAGCTTCCCGTTCTCAG-3’; *CREB1:* F 5’-GACCACTGATGGACAGCAGATC-3’ and R 5’-GAGGATGCCATAACAACTCCAGG-3’; *GNAS:* F 5’-CTCGCTGAGAAAGTCCTTGC-3’ and R 5’-CGAATGAAGTACTTGGCCCG-3’; *ARRB2*: F 5’-CTGACTACCTGAAGGACCGCAA-3’ and R 5’-GTGGCGATGAACAGGTCTTTGC-3’; *NR4A1:* F 5’-GGACAACGCTTCATGCCAGCAT-3’ and R 5’-CCTTGTTAGCCAGGCAGATGTAC-3’; *PCK1:* F 5’-CTGCCCAAGATCTTCCATGT-3’ and R 5’-CAGCACCCTGGAGTTCTCTC-3’; *NUP93:* F 5’-ACCGACAACCAGAGTGAAGTGG-3’ and R 5’-CGCCATAGTCTTCCAACAACTGC-3’; *PKMYT1:* F 5’-*TATGGGACAGCAGCGGATGTGT*-3’ and R 5’-AGAACGCAGCTCGGAAGACAGA-3’; *ADRB2:* F 5’-GATTTCAGGATTGCCTTCCA-3’ and F 5’-TATCCACTCTGCTCCCCTGT-3’. All transcript levels were normalized to the levels of the housekeeping gene *GAPDH*.

#### cAMP production

For pGLO sensor real-time measurement of cAMP production, sgRNA transduced cells seeded on 6-well plates at ~60-70% confluency were transfected with 900 ng pGLO-20F/Rluc for 24 h using Lipofectamine 2000 transfection reagent following manufacturer’s protocols. On the day of the experiment, cells were replated onto a 96-well plate in medium supplemented with 1.6 mmol D-luciferin and incubated for 40 min prior to conducting the assay. Hamamatsu FDSS/μCell with liquid handling was equilibrated at 37°C and used to add the drugs (1 μM isoproterenol, 10 μM NECA or 10 μM forskolin) and simultaneously image cAMP-driven luciferase activity in real time. At the end of the time course, cells were lysed in stop buffer (5 mM HEPES, 2% glycerol, 1 mM EDTA, 400 uM DTT, 0.2% Triton) supplemented with 2 μM coelenterazine, and all experimental cAMP measurements (firefly luciferase time course data) were normalized to the Renilla luciferase signal (expression control). The average normalized maximum values from the control samples for each batch tested was set to 100%, and all values are shown as % of this mean. For the competitive ELISA cAMP assay, cells transduced with sgRNAs were plated on 6-well plates and lysed 7 days post-infection by addition of 0.1 M HCl either in the absence of stimulation or after a 5-min treatment with 1 μM isoproterenol. Assay was performed using a commercial kit (Cayman Chemical Cat #581001) following manufacturer recommendations. All values were normalized to total protein concentration of the respective sample. The average isoproterenol treated control pmol/ml for each test batch was set to 100%, and all values are shown as % of this value.

#### Receptor trafficking

sgRNA transduced cells seeded on 6-well plates at ~60% confluency were transfected with 2 μg flag-tagged β2-AR for 48 hr using JetPrime Transfection reagent (Genesee, Cat #55134) following manufacturer’s protocols. To induce β2-AR internalization, cells were treated with 1 μM isoproterenol for 20 min. Then, cells were put on ice to stop trafficking, lifted and labeled with Alexa 647-conjugated M1 antibody. Untreated cells served as a control for total β2-AR cell surface expression. Flow cytometry analysis of 10,000 cells total per sample was carried out using a BD FACS Canto2 instrument, and Alexa-647 mean signal of the gated singlet population was used as a proxy for total number of surface receptors. Calculations were carried out for each cell line as follows: % internalized receptors = (total # surface receptors after 20 min isoproterenol)/(total # surface receptors pre-drug) × 100 %. The mean Alexa-647 control values for the batch was set to 100%, and all values are shown as % of this mean.

### Accession codes

Datasets from the CRISPRi screen were deposited on Gene Expression Omnibus under accession number GSE140745.

## Supplementary Figures

**Figure S1. A CREB reporter for flow cytometry. (A)** cAMP accumulation following β2-AR activation with isoproterenol or direct adenylyl cyclase stimulation with forskolin. Cells transfected with pGLO-20F were stimulated with indicated doses of isoproterenol (Iso) or with 10 μM forskolin. Arrow indicates the time point of drug treatment. The maximum luminescence from 10 μM forskolin was set to 100 and all isoproterenol values were normalized to that. Data plotted are from n = 3 per condition. **(B)** CRE-DD-GFP reporter accumulation was assessed in transiently transfected HEK293 cells in the presence of absence of Shield-1by flow cytometry. Cells were treated with 1 μM isoproterenol or 10 μM forskolin (FSK) in the presence or absence of the ProteoTuner stabilizing compound, Shield-1, for 4 h, and reporter expression was analyzed by flow cytometry. Data are mean GFP expression from GFP+-gated singlet cells (“% expressing cells”), n = 2 per condition. **(C)** A CRE-GFP/CRISPR clonal reporter cell line responds robustly to β2-AR stimulation by isoproterenol in a dose-dependent manner. Cells were treated with indicated doses of isoproterenol and 1 μM Shield-1 for 4 h, and reporter expression was analyzed by flow cytometry. Data plotted are from n = 3 per condition.

**Figure S2. Efficient CRISPR-dependent depletion of *TFRC* protein levels and inhibition of *ADRB2* receptor function in CRE-GFP/CRISPR cells. (A)** sgRNA against *TFRC* successfully depletes receptor expression. Cell-surface transferrin receptors were labeled with a PE-Cy7-conjugated antibody and expression levels were quantified by flow cytometry. Data are means from n = 2. **(B)** Knockdown of *ADRB2* by CRISPRi disrupts CRE-DD-GFP accumulation. In sgRNA-infected cells (1 NTC and 2 *ADRB2*-specific sgRNAs), cAMP signaling was activated either with 1 μM isoproterenol or 1 mM 8-Bromo-cAMP in the presence of 1 μM Shield-1 for 4 h, and CREB reporter expression was quantified by flow cytometry. Data are means from n = 2-5. In each flow cytometry experiment, 10,000 cells total were analyzed and gated for singlets. Error bars = ± s.e.m. ** = *p* ≤ 0.01 by unpaired two-tailed Student’s t-test, **** = *p* ≤ 0.0001 by one-way ANOVA test.

**Figure S3. Validation of candidate regulators identified in the “medium”/”low” fractions of the screen. (A)** Volcano plot of gene enrichment in the “medium” versus “low” reporter-expressing sorted fractions from all seven CRISPRi libraries. Fold enrichment is calculated as the log_2_ of the mean counts for 3/5 sgRNAs targeting a given gene, while *p* values are calculated based on the distribution of all 5/5 sgRNA targeting a gene relative to the NTCs in the library. Significant hits with “hit strength” FDR ≤ 10% are color-coded based on their phenotype, and *HRAS* and *MICAL3* (arrows) were selected for follow-up validation. Phenotype associated with *ADRB2* depletion is indicated for reference. **(B-C)** Depletion of *HRAS* **(B)** results in mild upregulation of reporter levels, while depletion of *MICAL3* **(C)** has no effect on isoproterenol-induced CRE-DD-GFP accumulation. Two *MICAL3* and three *HRAS* sgRNAs were selected, individually cloned and tested. Data shown are average from n = 2-4 per sgRNA. Error bars = ± s.e.m. * = *p* ≤ 0.05 by one-way ANOVA test.

**Figure S4. Effects of CRISPRi-dependent depletion of known regulators of cAMP signaling. (A)** Enrichment of *CREB1*-, *GNAS*-, and *ARRB2*-targeting sgRNAs in “high” vs “low” sorted population are plotted and color-coded based on the log_2_ enrichment of sgRNA counts as indicated in the legend. Negative control sgRNAs are colored grey. sgRNAs used for follow-up analyses are indicated. **(B)** No effect on reporter accumulation from silencing of *CREB1*, *GNAS* and *ARRB2* with individually cloned and transduced sgRNAs. Cells were treated with 1 μM Shield-1 alone (basal, ND = no drug) or with 1 μM isoproterenol in the presence of 1 μM Shield-1 (ISO) for 4 h, and analyzed by flow cytometry. The mean GFP signal of the two control sgRNAs (5443 and 5444) was set to 100% and used to normalize the mean GFP values for all sgRNAs from that batch. Data are mean from n = 3 (gene-specific sgRNAs) and n = 5-6 (NTCs). **(C)** mRNA depletion for *CREB1*, *GNAS* and *ARRB2*. Levels of each transcript in unstimulated cells were quantified by real-time PCR and normalized to *GAPDH*. Data are mean from n = 3. **(D)** CRISPRi-dependent depletion of beta-arrestin leads to persistent cAMP production upon stimulation with 1 μM isoproterenol measured with pGLO-RLuc assay. Data are mean from n = 3. **(E)** *ARRB2* is required for β2-AR internalization measured by flow cytometry of transiently transfected receptors in CRISPRi cells. Total of 10,000 cells were measured and gated for singlets. Data are mean from n = 3. **(E)** Silencing of beta-arrestin does not impact cAMP production stimulated by 10 μM forskolin measured with pGLO-RLuc assay. Data are mean from n = 3. **(G)** Depletion of beta-arrestin by CRISPRi and transcriptional upregulation of the known CREB targets, *NR4A1* (right) and *PCK1* (left). Data are *GAPDH-*normalized means from n = 3. Error bars = ± s.e.m. * = *p* ≤ 0.05 by one-way ANOVA test.

**Figure S5. Effects of validated high-confidence hits on basal and induced CRE-GFP accumulation.** Cells were treated with 1 μM Shield-1 alone (basal), with 10 μM NECA (A2-R) or 10 μM FSK in the presence of 1 μM Shield-1 for 4 h, and analyzed by flow cytometry. For all experiments, two sgRNAs per gene were selected based on screen phenotype, individually cloned and tested in batches. The mean GFP signal of the two control sgRNAs (5443 and 5444) for each batch was set to 100% and used to normalize the mean GFP values for all sgRNAs from that batch, as described in the “Materials and Methods” section. Effects of depleting select novel regulators of signaling verified in Fig. 3b on CRE-GFP reporter levels under basal **(A)**, A2-R-**(B)** or forskolin-induced **(C)** conditions. Data shown are mean from from n = 3-12 (gene-specific sgRNAs) and n = 40 (NTCs). Error bars = ± s.e.m. **** = *p* ≤ 0.0001; *** = *p* ≤ 0.001; ** = *p* ≤ 0.01; * = *p* ≤ 0.05 by one-way ANOVA test.

**Figure S6. Functional characterization of *NUP93* and *PKMYT1*. (A)** mRNA depletion of *NUP93* and *PKMYT1* by CRISPRi measured by qPCR and normalized to the levels of *GAPDH*. Data are mean from n = 4 (*PKMYT1*) and 6 (*NUP93*). **(B)** Effects of *NUP93* knockdown on basal and stimulated levels of a known endogenous CREB target, *NR4A1*. Cells were treated with 1 μM isoproterenol or 10 μM forskolin for 1 h, RNA was isolated, reverse transcribed and analyzed by qPCR. Data plotted are *GAPDH*-normalized mean levels from n = 3 (FSK) and 6 (basal, ISO). **(C)** *NUP93* knockdown diminishes cAMP production following 5-min stimulation with 1 μM isoproterenol measured by cAMP ELISA assay. All values were normalized to total protein per sample and shown as percent of averaged controls. Data are mean from n = 3-5. **(D)** *PKMYT1* knockdown does not have significant effects on bulk cAMP production following GPCR-dependent (1 μM isoproterenol, 10 μM NECA) and direct induction of adenylyl cyclase enzymes (10 μM FSK). cAMP accumulation was measured by pGLO luciferase assay, the maximum biosensor values were normalized to plasmid transfection, and are shown as percent of averaged controls. Data are mean from n = 9. **(E)** Effect of *NUP93* depletion on *ADRB2* levels quantified by RT-qPCR. Data plotted are *GAPDH-*normalized mean levels from n = 4. **(F)** Effects of *PKMYT1* depletion on trafficking of β2-AR analyzed by flow cytometry. Cells transiently transfected with flag-tagged β2-AR were either left untreated or were induced with 1 μM isoproterenol for 20 min, and cell-surface levels of flag-tagged receptor were measured by incubation with Alexa-647-M1 on ice and flow cytometry analysis. Average control values were set to 100% and used to normalize the data as described in the “Materials and Methods” section. Data are mean from n = 3 (*PKMYT1* sgRNAs) and 5 (NTCs). Error bars = ± s.e.m. *** = *p* ≤ 0.001; ** = *p* ≤ 0.01; * = *p* ≤ 0.05 by one-way ANOVA test.

## References

1 Hauser, A. S., Attwood, M. M., Rask-Andersen, M., Schioth, H. B. & Gloriam, D. E. Trends in GPCR drug discovery: new agents, targets and indications. Nat Rev Drug Discov 16, 829–842, doi:10.1038/nrd.2017.178 (2017).

2 Sharma, S. & Petsalaki, E. Application of CRISPR-Cas9 Based Genome-Wide Screening Approaches to Study Cellular Signalling Mechanisms. IntJ Mol Sci 19, doi:10.3390/ijms19040933 (2018).

3 Lohse, M. J., Engelhardt, S. & Eschenhagen, T. What is the role of beta-adrenergic signaling in heart failure? Circ Res 93, 896–906, doi:10.1161/01.RES.0000102042.83024.CA (2003).

4 Mutlu, G. M. & Factor, P. Alveolar epithelial beta2-adrenergic receptors. Am J Respir Cell Mol Biol 38, 127–134, doi:10.1165/rcmb.2007-0198TR (2008).

5 Kampmann, M. CRISPRi and CRISPRa Screens in Mammalian Cells for Precision Biology and Medicine. ACS Chem Biol 13, 406–416, doi:10.1021/acschembio.7b00657 (2018).

6 Maynard-Smith, L. A., Chen, L. C., Banaszynski, L. A., Ooi, A. G. & Wandless, T. J. A directed approach for engineering conditional protein stability using biologically silent small molecules. J Biol Chem 282, 24866–24872, doi:10.1074/jbc.M703902200 (2007).

7 Tsvetanova, N. G. & von Zastrow, M. Spatial encoding of cyclic AMP signaling specificity by GPCR endocytosis. Nature chemical biology 10, 1061–1065, doi:10.1038/nchembio.1665 (2014).

8 Barlow, C. A. et al. Protein kinase A-mediated CREB phosphorylation is an oxidant-induced survival pathway in alveolar type II cells. Apoptosis 13, 681–692, doi:10.1007/s10495-008-0203-z (2008).

9 Stern, C. M., Luoma, J. I., Meitzen, J. & Mermelstein, P. G. Corticotropin releasing factor-induced CREB activation in striatal neurons occurs via a novel Gbetagamma signaling pathway. PLoS One 6, e18114, doi:10.1371/journal.pone.0018114 (2011).

10 Xie, F. et al. Identification of a Potent Inhibitor of CREB-Mediated Gene Transcription with Efficacious in Vivo Anticancer Activity. J Med Chem 58, 5075–5087, doi:10.1021/acs.jmedchem.5b00468 (2015).

11 Tan, Y. et al. Selection and identification of transferrin receptor-specific peptides as recognition probes for cancer cells. Anal Bioanal Chem 410, 1071–1077, doi:10.1007/s00216-017-0664-4 (2018).

12 Fredericks, Z. L., Pitcher, J. A. & Lefkowitz, R. J. Identification of the G protein-coupled receptor kinase phosphorylation sites in the human beta2-adrenergic receptor. J Biol Chem 271, 13796–13803, doi:10.1074/jbc.271.23.13796 (1996).

13 Nobles, K. N. et al. Distinct phosphorylation sites on the beta(2)-adrenergic receptor establish a barcode that encodes differential functions of betaarrestin. Sci Signal 4, ra51, doi:10.1126/scisignal.2001707 (2011).

14 Violin, J. D. et al. beta2-adrenergic receptor signaling and desensitization elucidated by quantitative modeling of real time cAMP dynamics. The Journal of biological chemistry 283, 2949–2961, doi:10.1074/jbc.M707009200 (2008).

15 Gilbert, L. A. et al. Genome-Scale CRISPR-Mediated Control of Gene Repression and Activation. Cell 159, 647–661, doi:10.1016/j.cell.2014.09.029 (2014).

16 Thul, P. J. et al. A subcellular map of the human proteome. Science 356, doi:10.1126/science.aal3321 (2017).

17 Kampmann, M., Bassik, M. C. & Weissman, J. S. Functional genomics platform for pooled screening and generation of mammalian genetic interaction maps. Nat Protoc 9, 1825–1847, doi:10.1038/nprot.2014.103 (2014).

18 Kim, J. et al. Functional antagonism of different G protein-coupled receptor kinases for beta-arrestin-mediated angiotensin II receptor signaling. Proc Natl Acad Sci U S A 102, 1442–1447, doi:10.1073/pnas.0409532102 (2005).

19 Ren, X. R. et al. Different G protein-coupled receptor kinases govern G protein and beta-arrestin-mediated signaling of V2 vasopressin receptor. Proc Natl Acad Sci US A 102, 1448–1453, doi:10.1073/pnas.0409534102 (2005).

20 Chen, J. J. et al. Compromised function of the ESCRT pathway promotes endolysosomal escape of tau seeds and propagation of tau aggregation. J Biol Chem 294, 18952–18966, doi:10.1074/jbc.RA119.009432 (2019).

21 Ramkumar, P. et al. CRISPR-based screens uncover determinants of immunotherapy response in multiple myeloma. Blood Adv 4, 2899–2911, doi:10.1182/bloodadvances.2019001346 (2020).

22 Tian, R. et al. CRISPR Interference-Based Platform for Multimodal Genetic Screens in Human iPSC-Derived Neurons. Neuron 104, 239–255 e212, doi:10.1016/j.neuron.2019.07.014 (2019).

23 Ashburner, M. et al. Gene ontology: tool for the unification of biology. The Gene Ontology Consortium. Nat Genet 25, 25–29, doi:10.1038/75556 (2000).

24 Mi, H. et al. PANTHER version 11: expanded annotation data from Gene Ontology and Reactome pathways, and data analysis tool enhancements. Nucleic Acids Res 45, D183–D189, doi:10.1093/nar/gkw1138 (2017).

25 The Gene Ontology, C. The Gene Ontology Resource: 20 years and still GOing strong. Nucleic Acids Res 47, D330–D338, doi:10.1093/nar/gky1055 (2019).

26 Brown, C. R., Kennedy, C. J., Delmar, V. A., Forbes, D. J. & Silver, P. A. Global histone acetylation induces functional genomic reorganization at mammalian nuclear pore complexes. Genes Dev 22, 627–639, doi:10.1101/gad.1632708 (2008).

27 Chen, X. & Xu, L. Specific nucleoporin requirement for Smad nuclear translocation. Mol Cell Biol 30, 4022–4034, doi:10.1128/MCB.00124-10 (2010).

28 Ibarra, A., Benner, C., Tyagi, S., Cool, J. & Hetzer, M. W. Nucleoporin-mediated regulation of cell identity genes. Genes Dev 30, 2253–2258, doi:10.1101/gad.287417.116 (2016).

29 Labade, A. S., Karmodiya, K. & Sengupta, K. HOXA repression is mediated by nucleoporin Nup93 assisted by its interactors Nup188 and Nup205. Epigenetics Chromatin 9, 54, doi:10.1186/s13072-016-0106-0 (2016).

30 Liu, F., Stanton, J. J., Wu, Z. & Piwnica-Worms, H. The human Myt1 kinase preferentially phosphorylates Cdc2 on threonine 14 and localizes to the endoplasmic reticulum and Golgi complex. Mol Cell Biol 17, 571–583, doi:10.1128/mcb.17.2.571 (1997).

31 Liu, L. et al. PKMYT1 promoted the growth and motility of hepatocellular carcinoma cells by activating beta-catenin/TCF signaling. Exp Cell Res 358, 209–216, doi:10.1016/j.yexcr.2017.06.014 (2017).

32 Hanyaloglu, A. C. & von Zastrow, M. Regulation of GPCRs by endocytic membrane trafficking and its potential implications. Annu Rev Pharmacol Toxicol 48, 537–568, doi:10.1146/annurev.pharmtox.48.113006.094830 (2008).

33 Nigg, E. A., Schafer, G., Hilz, H. & Eppenberger, H. M. Cyclic-AMP-dependent protein kinase type II is associated with the Golgi complex and with centrosomes. Cell 41, 1039–1051, doi:10.1016/s0092-8674(85)80084-2 (1985).

34 Godbole, A., Lyga, S., Lohse, M. J. & Calebiro, D. Internalized TSH receptors en route to the TGN induce local Gs-protein signaling and gene transcription. Nat Commun 8, 443, doi:10.1038/s41467-017-00357-2 (2017).

35 Antoni, F. A., Wiegand, U. K., Black, J. & Simpson, J. Cellular localisation of adenylyl cyclase: a post-genome perspective. Neurochem Res 31, 287–295, doi:10.1007/s11064-005-9019-1 (2006).

36 Calebiro, D. et al. Persistent cAMP-signals triggered by internalized G-protein-coupled receptors. PLoS Biol 7, e1000172, doi:10.1371/journal.pbio.1000172 (2009).

37 Kotowski, S. J., Hopf, F. W., Seif, T., Bonci, A. & von Zastrow, M. Endocytosis promotes rapid dopaminergic signaling. Neuron 71, 278–290, doi:10.1016/j.neuron.2011.05.036 (2011).

38 Lazar, A. M. et al. G protein-regulated endocytic trafficking of adenylyl cyclase type 9. Elife 9, doi:10.7554/eLife.58039 (2020).

39 Steinberg, F. et al. A global analysis of SNX27-retromer assembly and cargo specificity reveals a function in glucose and metal ion transport. Nat Cell Biol 15, 461–471, doi:10.1038/ncb2721 (2013).

40 Horlbeck, M. A. et al. Mapping the Genetic Landscape of Human Cells. Cell 174, 953–967 e922, doi:10.1016/j.cell.2018.06.010 (2018).

41 Tsvetanova, N. G. et al. G Protein-Coupled Receptor Endocytosis Confers Uniformity in Responses to Chemically Distinct Ligands. Mol Pharmacol 91, 145–156, doi:10.1124/mol.116.106369 (2017).

42 Bingham, J., Sudarsanam, S. & Srinivasan, S. Profiling human phosphodiesterase genes and splice isoforms. Biochem Biophys Res Commun 350, 25–32, doi:10.1016/j.bbrc.2006.08.180 (2006).

43 Stelzer, G. et al. The GeneCards Suite: From Gene Data Mining to Disease Genome Sequence Analyses. Curr Protoc Bioinformatics 54, 1 30 31–31 30 33, doi:10.1002/cpbi.5 (2016).

44 Dugan, K. A., Wood, M. A. & Cole, M. D. TIP49, but not TRRAP, modulates c-Myc and E2F1 dependent apoptosis. Oncogene 21, 5835–5843, doi:10.1038/sj.onc.1205763 (2002).

45 Feng, Y., Lee, N. & Fearon, E. R. TIP49 regulates beta-catenin-mediated neoplastic transformation and T-cell factor target gene induction via effects on chromatin remodeling. Cancer Res 63, 8726–8734 (2003).

46 Giebler, H. A., Lemasson, I. & Nyborg, J. K. p53 recruitment of CREB binding protein mediated through phosphorylated CREB: a novel pathway of tumor suppressor regulation. Mol Cell Biol 20, 4849–4858, doi:10.1128/mcb.20.13.4849-4858.2000 (2000).

47 Van Orden, K., Giebler, H. A., Lemasson, I., Gonzales, M. & Nyborg, J. K. Binding of p53 to the KIX domain of CREB binding protein. A potential link to human T-cell leukemia virus, type I-associated leukemogenesis. J Biol Chem 274, 26321–26328, doi:10.1074/jbc.274.37.26321 (1999).

48 Gilbert, L. A. et al. CRISPR-mediated modular RNA-guided regulation of transcription in eukaryotes. Cell 154, 442–451, doi:10.1016/j.cell.2013.06.044 (2013).

49 Horlbeck, M. A. et al. Compact and highly active next-generation libraries for CRISPR-mediated gene repression and activation. Elife 5, doi:10.7554/eLife.19760 (2016).

50 Langmead, B., Trapnell, C., Pop, M. & Salzberg, S. L. Ultrafast and memory-efficient alignment of short DNA sequences to the human genome. Genome Biol 10, R25, doi:10.1186/gb-2009-10-3-r25 (2009).

