## Supplementary figures and images for "A high-throughput CRISPR interference screen for dissecting functional regulators of GPCR/cAMP signaling"

### Supplementary Figure 1

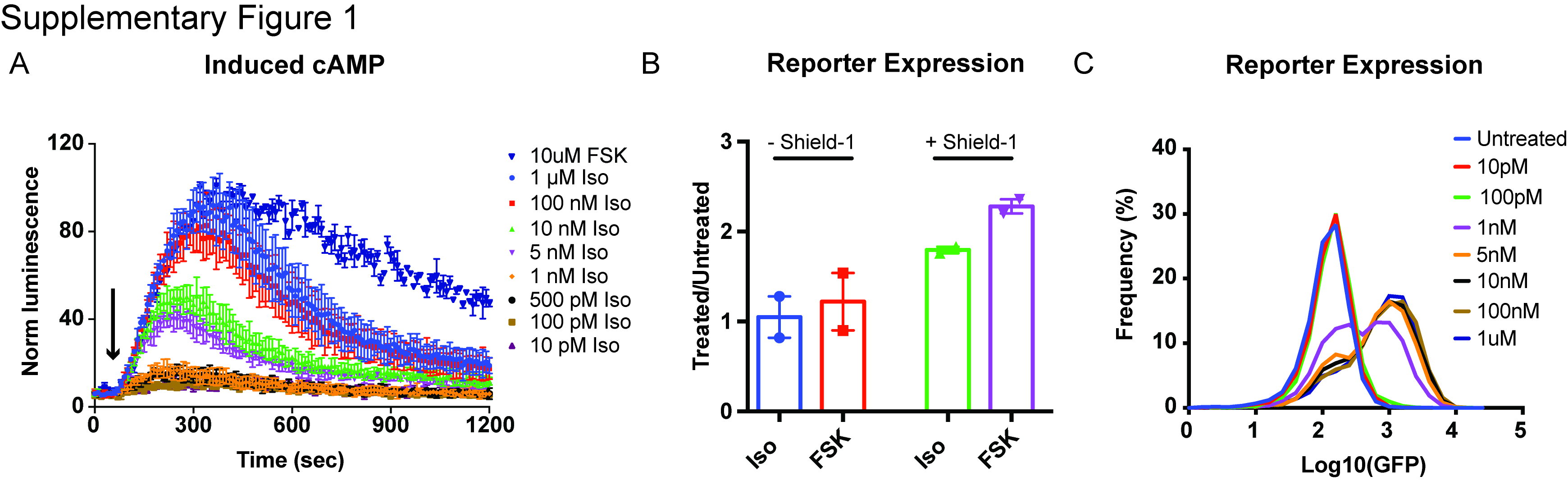

### Supplementary Figure 2

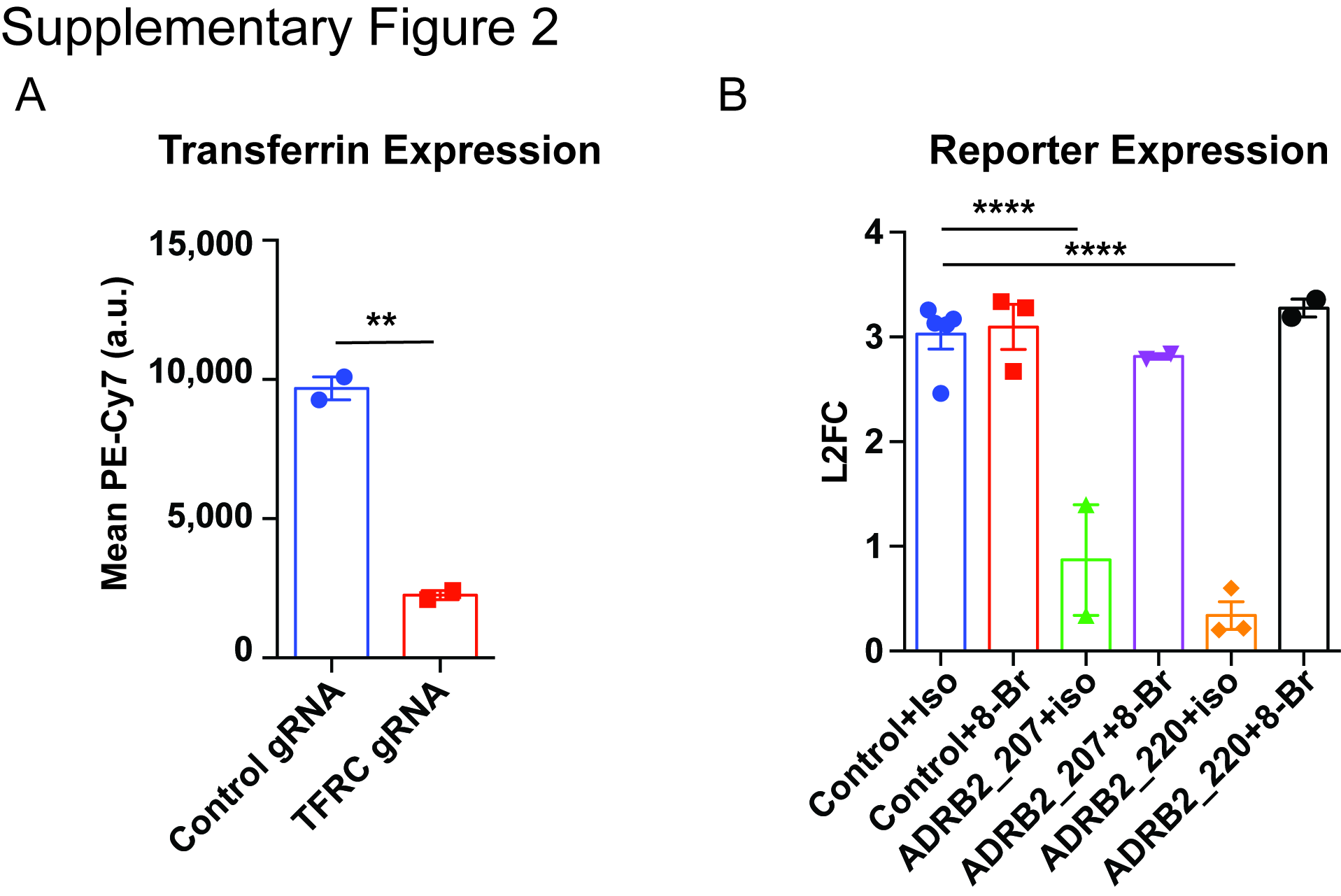

### Supplementary Figure 3

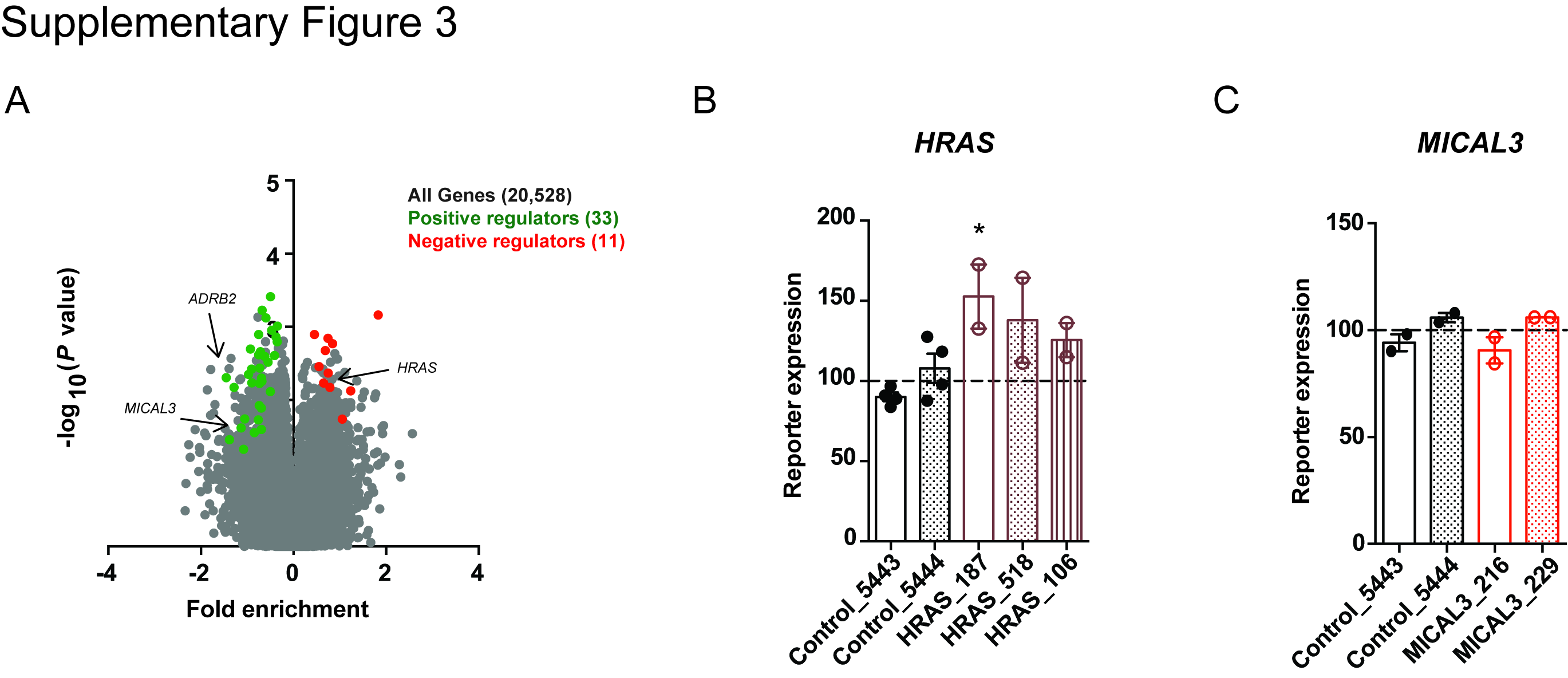

### Supplementary Figure 4

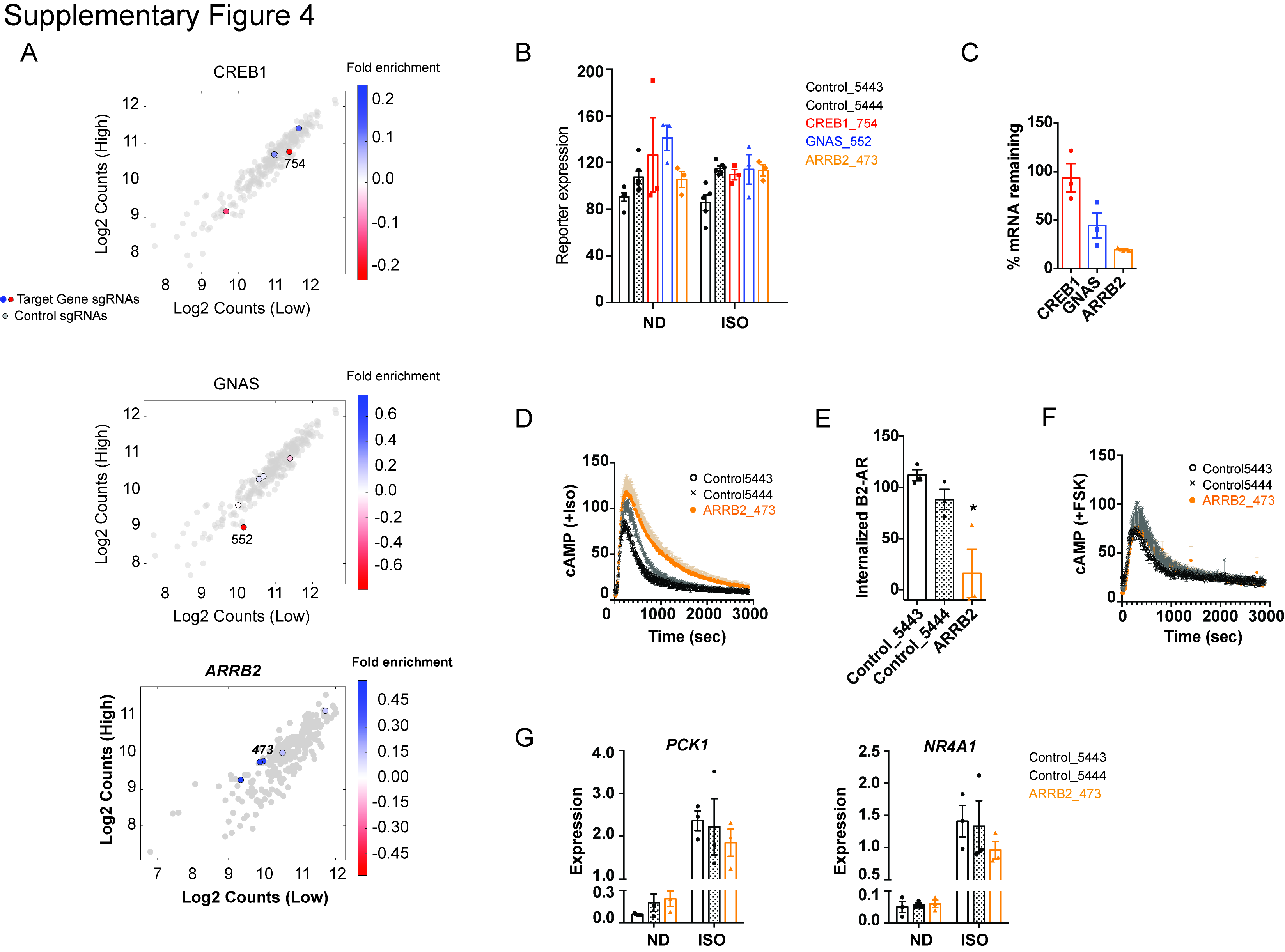

### Supplementary Figure 5

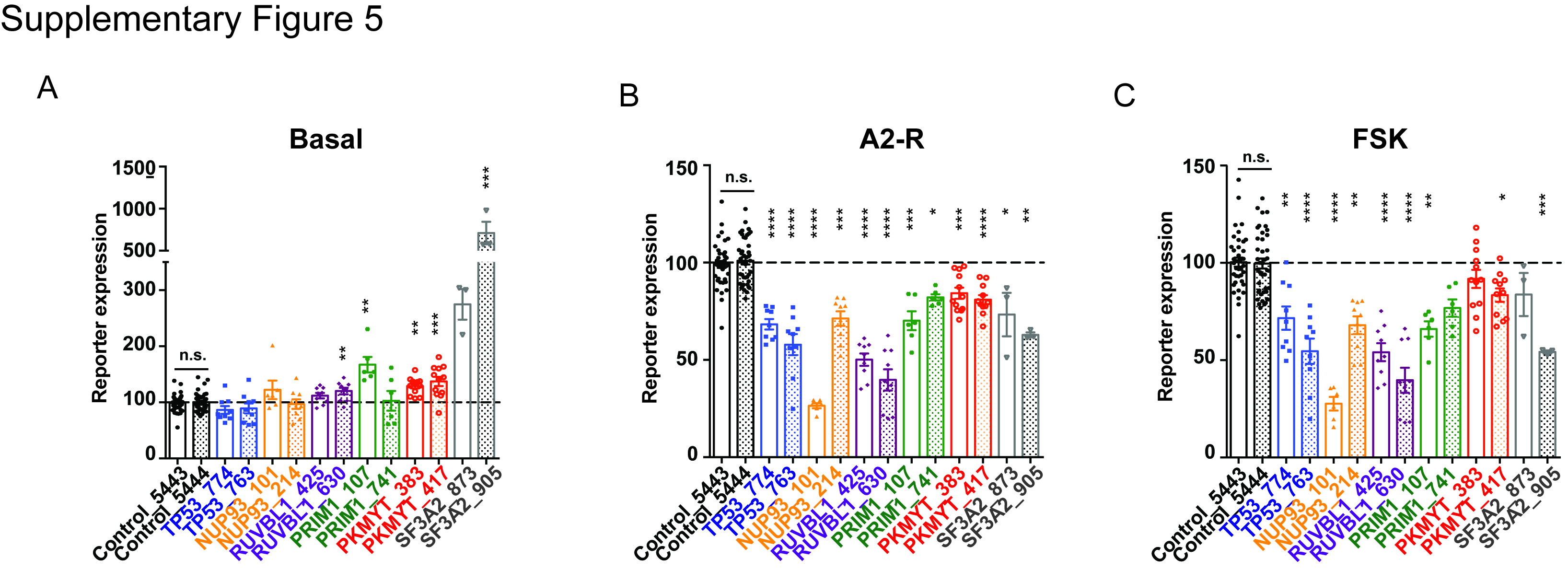

### Supplementary Figure 6

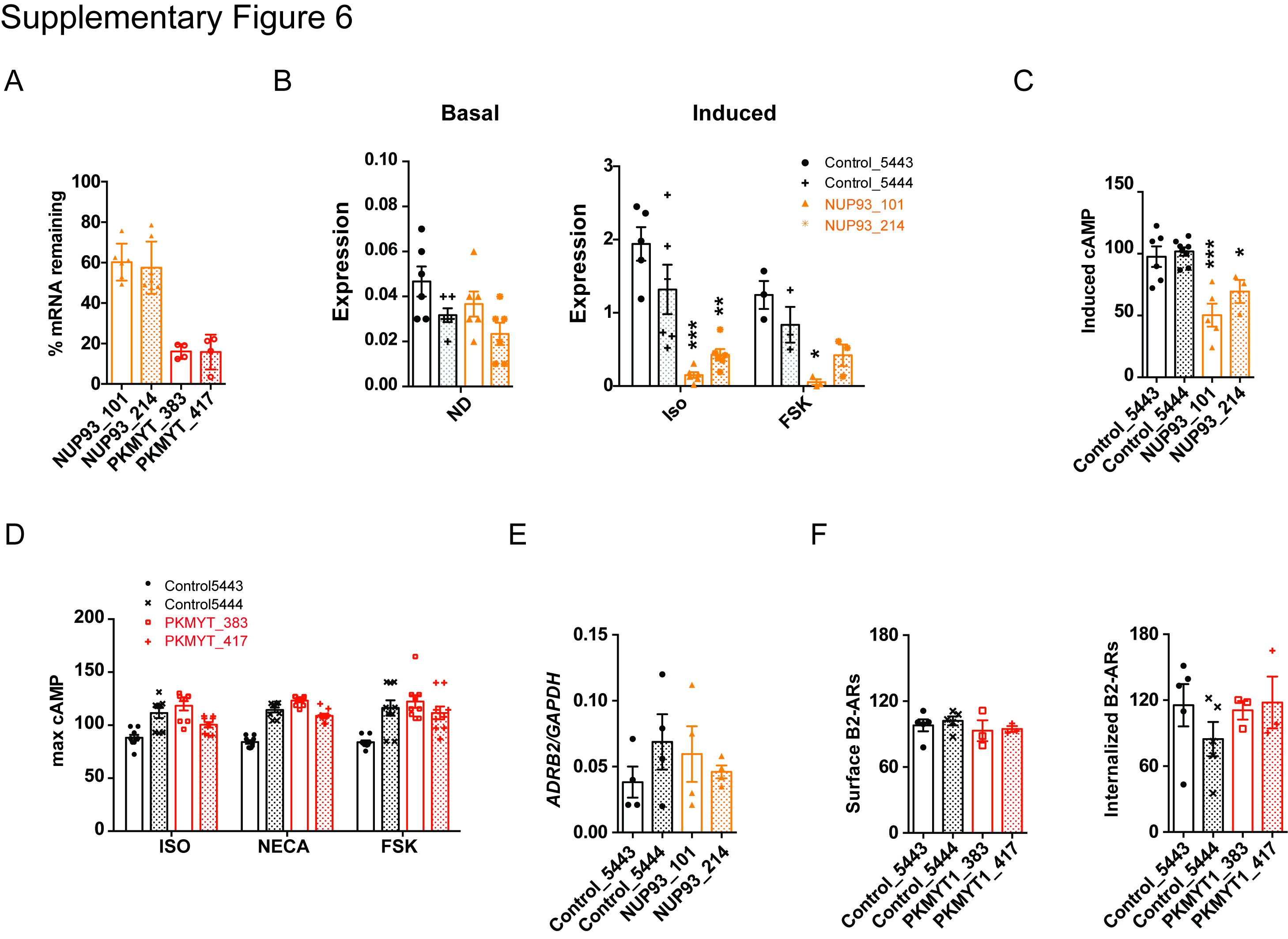
